# Inulin reduces *Ralstonia* invasion by promoting cooperation between *Lysinibacillus sphaericus* and *Pseudarthrobacter* spp.

**DOI:** 10.1101/2025.04.28.650914

**Authors:** Tao Yan, Ying Guan, Xinlan Mei, Ziru Zhang, Tianxiang Zhu, Yang Gao, Xiaofang Wang, Yangchun Xu, Qirong Shen, Zhong Wei, Tianjie Yang

## Abstract

In field production, beneficial microbial agents are typically applied along with key resources that beneficial microbes prefer to utilize. This practice aims to help the beneficial microbes colonize the field and exert their disease resistance functions. However, the mechanisms by which these key resources influence the colonization and disease resistance effects of beneficial microbes remain unknown. In this study, we found that inulin can enhance the growth capacity of *Lysinibacillus sphaericus* HR92 and its ability to inhibit *Ralstonia solanacearum*. Further investigation revealed that inulin can enhance strain HR92’s flagellar assembly, bacterial chemotaxis, fatty acid metabolism, and siderophore synthesis capabilities, ultimately improving strain HR92’s effectiveness in controlling bacterial wilt. Moreover, adding inulin to the soil can enrich *Pseudarthrobacter* strains. This genus enhances the expression of genes involved in the synthesis of the antibacterial substance HR92, and competes for nutrients with *Ralstonia solanacearum*, thereby further inhibiting the occurrence of bacterial wilt disease. Additionally, the combination of inulin and *Pseudarthrobacter* strains upregulates the valine and leucine pathway genes in strain HR92. These genes play a crucial role in the synthesis of surfactin and the antimicrobial VOC 3-methyl-1-butanol. We have demonstrated that the addition of key resources not only enhances the control efficacy of beneficial bacteria against soil-borne diseases but also enriches the “helpers” of beneficial bacteria in soil.

**Importance:** *Ralstonia solanacearum* is a plant pathogen that can cause bacterial wilt in several important crops. Given the limitations of beneficial bacterial inoculation, we propose a strategy to enhance the biocontrol efficacy of beneficial microorganisms by utilizing key resources. In this study, we demonstrate that key resources inulin can improve the biological control effects of beneficial bacteria, thereby maintaining plant health. Furthermore, we found that increasing the concentration of inulin altered the diversity of the rhizosphere microbiota and specifically enriched the abundance of *Pseudarthrobacter*, which compete with pathogens for nutrients on one hand and, on the other hand, synergize with inulin to enhance the biocontrol efficacy of the beneficial bacterium HR92. Together, our study reveals that key resources can enhance the biocontrol efficacy of beneficial bacteria and enrich potential beneficial bacteria in resident microbial communities, further reducing tomato diseases.

## Introduction

Soil-borne plant pathogens cause major economic losses to agricultural production due to lack of efficient control methods (1–5). In recent years, researchers have increasingly explored the use of biological control methods, which are ecologically friendly and exhibit specificity in pathogen suppression (6–9). Beneficial microbes can prevent pathogenic invasion of plant root systems through various mechanisms, they can secrete antagonistic substances that directly inhibit the growth of pathogens or their “helpers” (10), preventing colonization or invasion of plant roots. Some beneficial bacteria compete with pathogens for rhizosphere resources, thereby suppressing pathogen growth through nutrient competition. Moreover, introduced beneficial microbes can stimulate soil native resident microbiomes (11), engaging in metabolic interactions that fortify the rhizosphere, creating a more resilient barrier that pathogens find difficult to breach (10, 12–15). Studies have shown that *Lysinibacillus sphaericus* is an effective biocontrol agent against bacterial wilt in tomatoes (16–19). Specifically, *L. sphaericus* inhibits the growth of *R. solanacearum* through the secretion of volatile organic compounds (VOCs) such as dicarboxylic esters and 3-methyl-1-butanol (20), as well as cyclic lipopeptides like surfactin and bacitracin (21–23).

Due to the limited availability of nutrients in the rhizosphere and the complex conditions of the field environment (24, 25), the inoculated beneficial bacteria often lack a competitive edge over soil native resident microbiomes, preventing them from efficiently colonizing the rhizosphere and exhibiting their disease-suppressive functions (26–28). Recent studies have proposed providing nutritional carriers for inoculated beneficial bacteria to enhance their biocontrol efficacy in the rhizosphere (29–34). For instance, adding organic fertilizers, biochar, or amino acid-based fertilizers along with the beneficial microbes has been shown to improve their survival in the rhizosphere. However, these organic carriers are generally utilized by a wide range of soil microorganisms, lacking specificity, and cannot be exclusively exploited by beneficial bacteria. In recent years, research in the field of gut microbiota has suggested adding resources preferentially utilized by beneficial bacteria along with their introduction to target and enhance their colonization in the gut (35, 36). Due to the analogous roles of the rhizosphere and the gut in host immunity and ecological function, we hypothesize that the application of targeted resources capable of selectively enhancing the colonization efficiency of beneficial bacteria in the rhizosphere will promote their stable establishment in the root environment (37–39), ultimately improving their biocontrol performance. Soil microorganisms in the rhizosphere are often strongly influenced by plant root exudates, with sugars in these exudates playing a crucial role in recruiting beneficial bacteria and maintaining plant health (40–42). Thus, we focused on soluble sugars from root exudates to identify a compound capable of selectively enhancing the colonization capacity of *Lysinibacillus sphaericus*. Through this approach, we aim to explore how such a resource influences the biocontrol potential of beneficial bacteria within the complex rhizosphere environment.

Unlocking the potential of specific resources to enhance the targeted colonization of beneficial bacteria in the rhizosphere, while understanding their effects on soil native microbial communities, is crucial for managing soil-borne diseases in agricultural production. We utilized *L. sphaericus* HR92 along with inulin, which has been shown to boost both the growth and antibacterial activity of strain HR92. By inoculating the beneficial bacteria and varying concentrations of inulin to the tomato roots, we observed the colonization potential of strain HR92 and its impact on the assembly of bacterial communities in the tomato rhizosphere. Our findings reveal that inulin not only directly enhances strain HR92 colonization and disease suppression capabilities but also enriches specific soil native resident microbiomes. These native microbes, on one hand, exhibit inhibitory effects on pathogens and, on the other, enhance the disease resistance of beneficial bacteria. Finally, we used transcriptomics to determine the influence of inulin and native soil microbes on strain HR92.

## Materials and methods

### Strains and growth conditions

The plant pathogen used in this study was *Ralstonia solanacearum* QL-Rs1115 (GenBank: GU390462) (43), and the beneficial bacterium was *Lysinibacillus sphaericus* HR92 (GenBank: JF700465.1) (19). The strain was routinely grown in nutrient broth medium at 30 ℃ with shaking at 170 rpm in NB medium (nutrient broth, 10.0 g glucose L^−1^; 5.0 g peptone L^−1^, 0.5 g yeast extract L^−1^, and 3.0 g beef extract L^−1^, pH 7.0).

In *vitro* co-culture experiments, we used strain *R. solanacearum* QL-RFP, that QL-Rs1115 was tagged with the stable pYC12-mCherry plasmid (excitation wavelength: 587 nm, absorption wavelength: 610 nm) (14). The tagged plasmid had no significant impact on growth and virulence of the pathogen *R. solanacearum* (14). The strain *L. sphaericus* HR92 carries green fluorescence plasmid (excitation wavelength: 488 nm, absorption wavelength: 510 nm). The pathogen cells were harvested by centrifugation (6000 rpm, 10 min), washed twice with sterile saline solution (0.9% NaCl), and diluted to a density of 10^8^ colony-forming units (CFU) ml^−1^ based on the optical density 600 (OD_600_ = 0.5) before all the experiments.

### Effects of various sugars on the growth of *Ralstonia solanacearum* and HR92

The bacterial strain HR92, and *R. solanacearum* QL-Rs1115 were grown overnight in NB medium at 30 °C with shaking at 170 rpm. Cells were washed three times with 0.85% NaCl and resuspended to an OD_600_ of 0.5 (approximate 10^7^ CFU mL^-1^). Bacteria was cultured in a 96-well microtiter plate. Each well contained a total of 200 μL culture liquid: 178 μL NB medium, 20 μL of 1mM sugars solution (final concentration 0.1mM), and 2 μL bacterial suspension. We set treatment without sugars (the same volume sterile water instead) as control. The microtiter plate was incubated at 30 ℃ with agitation of 170 rpm. Bacterial growth (OD_600_) was measured after 48 h by using a SpectraMax M5 spectrophotometer (Molecular Devices, Sunnyvale, CA, USA). All treatments were performed with three replicates.

### Effect of various sugars on the ability of strain HR92 to inhibit *R. solanacearum*

We selected sugars that did not directly promote the growth of *R. solanacearum* for subsequent experiments. The bacterial strains HR92 and R. solanacearum QL-RFP were grown overnight in NB medium at 30 °C with shaking at 170 rpm. Cells were washed three times with 0.85% NaCl and resuspended to an OD600 of 0.5 (approximately 10⁷ CFU mL⁻¹). The bacteria were cultured in 96-well microtiter plates. Each well contained a total of 200 μL of culture: 176 μL NB medium, 20 μL of 1 mM sugar solution (final concentration 0.1 mM), and 2 μL each of HR92 and QL-RFP. We set treatment without sugars (the same volume sterile water instead) as control. The microtiter plates were incubated at 30 °C with shaking at 170 rpm. Bacterial growth (OD_600_) was measured after 48 hours using a SpectraMax M5 spectrophotometer (Molecular Devices, Sunnyvale, CA, USA). In addition, red fluorescence of QL-RFP (excitation wavelength: 587 nm; emission wavelength: 610 nm) was measured, and the biomass of *R. solanacearum* under co-culture with HR92 and sugars was calculated using log (RFU/OD_600_). All treatments were performed with three replicates.

### Greenhouse experiment setup and sample collection

Surface-sterilized tomato seeds (*Solanum lycopersicum,* cultivar “Hong’ai”) were germinated on water-agar plates for 3 days before sowing into 72-well trays filled with growth substrate (50 g substrate per well; commercially available from Jiangsu-Xingnong Substrate Technology Co., Ltd). At the three-leaf stage (17 days after sowing), a subset of tomato plants of similar size was selected and transplanted into seedling cups containing natural soil (200 g per cell). Seedlings were grown under controlled greenhouse conditions (25-30 ℃, 16/8 h day/night cycle, 200 μmol m^−2^ s^−1^ light density, and 70% relative humidity).

### Part1: Inulin enhances the disease-suppressive effect of HR92

Seven days after transplantation, strain HR92 and inulin were inoculated together into the rhizosphere soil of each plant. The final density of strain HR92 was 10^8^ CFU g^−1^ soil. The final concentration of inulin was adjusted according to each treatment. Seven days after co-inoculation of strain HR92 and inulin, the pathogen *R. solanacearum* QL-Rs1115 was inoculated into the rhizosphere soil of each plant at a final density of 10^6^ CFU g^−1^ soil. The greenhouse experiment comprised four treatments, including only pathogen, only HR92, HR92 with 0.1% inulin (1 g kg^-1^ soil), and HR92 with 0.3% inulin (3 g kg^-1^ soil) (Fig. S1). Each treatment had four biological replicates, with each replicate consisting of five individual tomato plants, resulting in a total of 80 plants (20 plants per treatment x 4 treatments). Tomato plants regularly watered with sterile water, and their positions were randomized every two days. Seven days after inoculation with *R. solanacearum*, the tomato plants began to exhibit disease symptoms. Disease index was recorded seven days after transplantation on a scale of 0 to 4 (0, no signs of wilting; 1, 1%–25% lead area wilted; 2, 26%–50% leaf area wilted; 3, 51%–75% leaf area wilted; and 4, 76%–100% leaf area wilted), and was calculated using the formula: DS = ∑ (number of diseased plants with given index × DI)/ (total number of plants infected × highest DI) (44). Soil samples from the rhizosphere of both healthy and diseased plants were collected and stored at −80 ℃.

### Part II: P265, P397, and inulin enhance the disease-suppressive effect of HR92

In a separate set of experiments with a similar design, we further evaluated the effects of two key bacterial strains (P265 and P397) in enhancing the biocontrol effect of HR92. Seven days after transplantation, strains P265, P397, and HR92 were co-inoculated into the rhizosphere together with inulin. The total bacterial density was adjusted to 10⁸ CFU g⁻¹ soil. Seven days after this co-inoculation, *R. solanacearum* QL-Rs1115 was introduced at 10⁶ CFU g⁻¹ soil. Treatments and replication were conducted as in Part I, with appropriate controls included for comparison. Each treatment had four biological replicates, with each replicate consisting of five individual tomato plants, resulting in a total of 140 plants (20 plants per treatment x 7 treatments).

### DNA extraction and 16S rRNA gene amplicon sequencing

Genomic DNA was extracted from 0.5 g of soil using the PowerSoil^®^ DNA Isolation Kit (MoBio, Carlsbad, CA, USA) following the manufacturer’s instructions. The purity and concentration of DNA were determined using a NanoDrop ND-1000 spectrophotometer (Thermo Fisher Scientific, Carlsbad, CA, USA) and a Qubit 2.0 fluorometer (Thermo Fisher Scientific). The V4 region of the bacterial 16S rRNA gene was amplified with primers 563F and 802R (Table S3) (45). A two-step dual indexing strategy for Illumina MiSeq (Illumina, San Diego, CA, USA) sequencing was employed. The PCR thermocycler program included an initial denaturation temperature of 95 ℃ for 2 min, then 25 cycles of 95 ℃ for 30 s, 55 ℃ for 30 s, and 72 ℃ for 30 s, and a final elongation step of 72 ℃ for 5 min. The PCR amplification was confirmed by gel electrophoresis on 2% agarose gels. PCR products were purified using AxyPrep PCR Clean-up Kit (Axygen Biosciences, CA, USA). The following library construction and Illumina MiSeq sequencing (2 × 300 bp) were performed by Shanghai Biozeron Biological Technology (Shanghai, China).

Raw amplicon reads were processed using the DADA2 pipeline v.1.14.1 (46). In brief, reads were quality checked and primers were removed using trimLeft in the filter AndTrim function. According to the sequence quality, the 16S rRNA gene reads were filtered using default parameters (confidence level at 80%). Chimeras were removed after merging denoised pair-end sequences. Each unique amplicon sequence variant (ASV) was assigned to taxa according to SILVA database v.138.1 (47) for the 16S rRNA gene. For bacteria, non-bacterial ASVs including chloroplasts and mitochondrial reads were removed.

### Real-time quantitative PCR analysis

Quantification of strain HR92 (19) and *R. solanacearum* (48) was performed by qPCR using the qTOWER Real-Time PCR system (Analytik jena, Jena, Germany). The specific primers utilized are listed in Table S3. Twenty-microliter reactions were containing 10 μL of 2X TB Green Premix Ex Taq (TaKaRa, Beijing, China), 2 μL of DNA, 0.8 μL of each primer (10 μM), 0.4 μL of ROX Reference Dye (TaKaRa, Beijing, China) and 6 μL of Nuclease-free water. Thermal cycling conditions were as follows: 95 ℃ for 3 min, followed by 40 cycles of 95 ℃ for 30 s, 50 ℃ for 30 s, and 72 ℃ for 30 s. Standard curves were used to calculate the gene copy numbers of the target group for each reaction. Each sample was run in triplicate, and results were expressed as log10 values of the gene copy numbers per gram of soil.

### Screening of Key bacteria

Bacteria were isolated from the rhizosphere of five healthy tomato plants co-treated with HR92 and inulin. Soil samples (1 g) were suspended in sterile water, vortexed, and serially diluted before plating. One hundred microliters of serial dilutions (10^−4^-10^−7^) were plated onto 1/3 TSA plates (Tryptic Soy Agar, tryptone 5 g L^−1^, soy peptone 1.65 g L^−1^, NaCl 1.65 g L^−1^ and agar 15 g L^−1^). Plates were incubated at 28 ℃ until single colonies existed on the medium.

All bacterial isolates were identified based on 16S rRNA gene sequence using universal primers 27F and 1492R (Table S3) (49). PCR products were sequenced by Sangon Biotech (Shanghai Co., Ltd, China). 16S rRNA gene sequencing results were blasted against the NCBI database for homologous sequences.

### Effect of inulin on bacterial growth

The bacterial strain HR92, P265 (*Pseudarthrobacter siccitolerans*), and P397 (*Pseudarthrobacter oxydans*), and *R. solanacearum* QL-Rs1115 were grown overnight in NB medium at 30 ℃ with shaking at 170 rpm. Cells were washed three times with 0.85% NaCl and resuspended to an OD_600_ of 0.5 (approximate 10^7^ CFU mL^-1^). Bacteria was cultured in a 96-well microtiter plate. Each well contained a total of 200 μL culture liquid: 178 μL NB medium, 20 μL 3% inulin (final concentration 0.3%), and 2 μL bacterial suspension. We set treatment without inulin (the same volume sterile water instead) as control. The microtiter plate was incubated at 30 ℃ with agitation of 170 rpm. Bacterial growth (OD_600_) was measured after 48 h by using a SpectraMax M5 spectrophotometer (Molecular Devices, Sunnyvale, CA, USA). All treatments were performed with four replicates.

### Effects of supernatants from HR92, P265, or P397 on *R. solanacearum* growth induced by inulin

After measuring bacterial growth, we obtained bacterial supernatant induced by inulin (**supernatant a**) by centrifuging bacterial suspension at 3000 rpm for 15 min with a 96-well filter membrane plate. 2 μL of *R. solanacearum* QL-Rs1115 suspension (OD_600_ = 0.5) were added to a 96-well microtiter plate with each well containing 178 μl of NB medium and 20 μl of **supernatant a**. We set a treatment without supernatant (replaced with the same volume of sterile water) as the control. The plate was incubated at 30 ℃ with agitation of 170 rpm. Ralstonia growth (OD_600_) was measured after 48 h by using a SpectraMax M5 spectrophotometer (Molecular Devices, Sunnyvale, CA, USA). All treatments were performed with five replicates.

### Effect of inulin on the ability of strain HR92, P265, or P397 to inhibit *R. solanacearum* under co-culture conditions

The bacterial strain HR92, P265, P397 and *R. solanacearum* QL-RFP were grown overnight in NB medium at 30 ℃ with shaking at 170 rpm. Cells were washed three times with 0.85% NaCl and resuspended to an OD_600_ of 0.5 (approximate 10^7^ CFU mL^-1^). Bacteria was cultured in a 96-well microtiter plate. Each well contained a total of 200 μL culture liquid: 176 μL NB medium, 20 μL 3% inulin (final concentration 0.3%), 2 μL QL-RFP, and 2 μL of one of the following strains: P265, P397, or HR92. We set treatment without strain and inulin (the same volume sterile water instead) as control. The microtiter plate was incubated at 30 ℃ with agitation of 170 rpm. Bacterial growth (OD_600_) was measured after 48 h by using a SpectraMax M5 spectrophotometer (Molecular Devices, Sunnyvale, CA, USA). In addition, red fluorescence of QL-RFP (excitation wavelength: 587 nm; emission wavelength: 610 nm) was measured, and the biomass of *R. solanacearum* under co-culture with HR92 and sugars was calculated using log(RFU/OD_600_). All treatments were performed with four replicates.

### Carbon resource utilization patterns of the bacterial isolates

The resource utilization patterns of QL-RS1115, HR92, P265 or P397 were assessed across 74 distinct carbon resources (Table S2), including amino acids, organic acids, and sugars commonly found in tomato root exudates. The bacterial strains were grown overnight in NB medium at 30 ℃ with shaking at 170 rpm. Cells were washed three times with 0.85% NaCl and resuspended to an OD_600_ of 0.5 (approximate 10^7^ CFU mL^-1^). Then, 5 μL of each strain was added to a 384-well microtiter plate containing 40 μL of OS minimal medium (50) supplemented with 5 μL of 1 mM of different carbon sources (14). Plates were incubated at 30 ℃ with agitation at 170 rpm, and bacteria growth was measured after 48 h using a SpectraMax M5 spectrophotometer (Molecular Devices, Sunnyvale, CA, USA). We set treatment without carbon resource (the same volume sterile water instead) as control. T-tests were conducted to compare the biomass of bacteria under each resource condition with the biomass under the control condition. A significance level of *p* < 0.05 indicates that the bacteria can utilize the respective resource. Utilization efficiency was calculated using the formula: (biomass under resource condition - control)/ control x 100. Utilization efficiency was categorized into different levels: below 20% was marked as *, 20%-40% as **, 40%-60% as ***, and above 60% as ****. All treatments were performed with four replicates.

### Effects of *Pseudarthrobacter* and HR92 combination on *R. solanacearum* growth induced by inulin

Strains HR92, P265, P397, and *Ralstonia solanacearum* QL-Rs1115 were grown overnight in NB medium at 30 ℃ with shaking at 170 rpm. After incubation, the cells were washed three times with 0.85% NaCl and resuspended to an OD600 of 0.5 (approximately 10⁷ CFU/mL). The bacteria were inoculated into a 96-well microtiter plate, with each well containing a total volume of 200 μL: 176 μL NB medium, 20 μL 3% inulin (final concentration 0.3%), and 2 μL of HR92, P265, or P397 suspension. Wells without P265 or P397 (replaced with the same volume of sterile water) were used as controls. The microtiter plate was incubated at 30 °C with shaking at 170 rpm for 48 hours. After incubation, the bacterial suspension was centrifuged at 3000 rpm for 15 minutes using a 96-well filter membrane plate to obtain the bacterial supernatant (**supernatant b**).Then, 2 μL of *R. solanacearum* QL-Rs1115 suspension (OD600 = 0.5) was added to each well of a new 96-well microtiter plate containing 178 μL of NB medium and 20 μL of **supernatant b**. Wells without P265 or P397 (replaced with the same volume of sterile water, only HR92 was included) were used as controls for **supernatant b**. The plate was incubated at 30 ℃ with shaking at 170 rpm. After 48 hours, the growth of *R. solanacearum* (OD600) was measured using a SpectraMax M5 spectrophotometer (Molecular Devices, Sunnyvale, CA, USA). All treatments were performed with four replicates.

### Effect of *Pseudarthrobacter* spp. on the ability of strain HR92 to inhibit *R. solanacearum*

This experiment followed a three-step workflow that was similar to previous inhibition assays. Initially, 2 μL of bacterial suspension (OD_600_ = 0.5) of strain P265 or P397 were incubated in 198 μL of NB medium at 30 ℃ with agitation at 170 rpm for 48 h. The resulting bacterial suspension was transferred to a 96-well filter membrane plate and centrifuged at 3000 rpm for 15 min to collect the supernatant (**supernatant c-1**). Next, 2 μL of bacterial suspension (OD_600_ = 0.5) of strain HR92 was incubated with 178 μL of NB medium and 20 μL of **supernatant c-1** under same conditions for 48 h. The bacterial suspension was then centrifuged, and the supernatant (**supernatant c-2**) was collected. Finally, 2 μL of *R. solanacearum* suspension (OD_600_ = 0.5) was incubated with 178 μL of NB medium and 20 μL of **supernatant c-2** at 30 ℃ with agitation at 170 rpm for 48 h. We set treatment without **supernatant c-2** (the same volume sterile water instead) as control. Bacterial growth (OD_600_) was measured using a SpectraMax M5 spectrophotometer (Molecular Devices, Sunnyvale, CA, USA). The effect of *Pseudarthrobacter* spp. on the ability of strain HR92 to inhibit *R. solanacearum* was analyzed by examining the biomass of *R. solanacearum*. All treatments were performed with four replicates.

### Effect of strain HR92 on the ability of *Pseudarthrobacter* spp. to inhibit *R. solanacearum*

This experiment followed a three-step workflow similarly to the previous inhibition assays. Initially, 2 μL of strain HR92 suspension (OD_600_ = 0.5) were incubated with 198 μL of NB medium at 30 ℃ with agitation at 170 rpm for 48 h, and the supernatant (**supernatant d-1**) was collected after centrifugation. Next, 2 μL of *Pseudarthrobacter* strains (strain P265 or P397) suspension (OD_600_ = 0.5) was incubated with 178 μL of NB medium and 20 μL of **supernatant d-1** under similar conditions for 48 h, followed by centrifugation to collect the supernatant (**supernatant d-2**). Finally, 2 μL of *R. solanacearum* suspension (OD_600_ = 0.5) was incubated with 178 μL of NB medium and 20 μL of **supernatant d-2** at 30 ℃ with agitation at 170 rpm for 48 h. We set treatment without **supernatant d-2** (the same volume sterile water instead) as control. Bacterial growth (OD_600_) was measured using a SpectraMax M5 spectrophotometer (Molecular Devices, Sunnyvale, CA, USA). The effect of strain HR92 on the ability of *Pseudarthrobacter* spp. to inhibit *R. solanacearum* was analyzed by examining the biomass of *R. solanacearum*. All treatments were performed with four replicates.

### RNA extraction and RNA-seq

Total RNA was extracted from the bacterial cells using TRIzol^®^ Reagent according the manufacturer’s instructions (Invitrogen, CA, USA). Then RNA quality was determined using 2100 Bioanalyser (Agilent, CA, USA) and quantified using the ND-2000 (NanoDrop Technologies, DE, USA). RNA-seq strand-specific libraries were prepared following TruSeq RNA sample preparation Kit from Illumina (San Diego, CA), using 5μg of total RNA. Shortly, rRNA removal by RiboZero rRNA removal kit (Epicenter, Wisconsin, USA), fragmented using fragmentation buffer cDNA synthesis, end repair, A-base addition and ligation of the Illumina-indexed adaptors were performed according to Illumina’s protocol. Libraries were then size selected for cDNA target fragments of 200–300 bp on 2% Low Range Ultra Agarose followed by PCR amplified using Phusion DNA polymerase (New England Biolabs, Massachusetts, USA) for 15 PCR cycles. After quantified by TBS380, paired-end libraries were sequenced by Illumina NovaSeq 6000 sequencing (2 × 150bp, Shanghai BIOZERON Co., Ltd).

The raw paired end reads were trimmed and quality controlled by Trimmomatic with parameters (SLIDINGWINDOW:4:15 MINLEN:75) (version 0.36 http://www.usadellab.org/cms/uploads/supplementary/Trimmomatic). Then clean reads were separately aligned to reference genome with orientation mode using Rockhopper (http://cs.wellesley.edu/~btjaden/Rockhopper/) software. Rockhopper was a comprehensive and user-friendly system for computational analysis of bacterial RNA-seq data. As input, Rockhopper takes RNA sequencing reads generated by high-throughput sequencing technology. This software was used to calculate gene expression levels with default parameters.

### Differential expression analysis and functional enrichment

To identify differential expression genes (DEGs) between the two different samples, the expression level for each transcript was calculated using the fragments per kilobase of read per million mapped reads (RPKM) method. edgeR (51) was used for differential expression analysis. The DEGs between two samples were selected using the following criteria: the logarithmic of fold change was greater than 2 and the false discovery rate (FDR) should be less than 0.05. To understand the functions of the differential expressed genes, Gene Ontology (GO) functional enrichment and Kyoto Encyclopedia and Genomes (KEGG) pathway analysis were carried out by Goatools (52) and KOBAS (53) respectively. DEGs were significantly enriched in GO terms and metabolic pathways when their Bonferroni-corrected P-value was less than 0.05.

### Effect of 3-methyl-1-butanol on the growth of pathogen *R. solanacearum*

In order to investigate the effect of key microbial metabolite on pathogen growth, we used standard chemical compound of 3-methyl-1-butanol and *R. solanacearum* to conduct a plate confrontation assay. The experiment was conducted using divided Petri dishes, with a 1 cm diameter filter paper was placed on one side and NA medium on the other. 3-methyl-1-butanol at concentrations ranging from 100% purity to dilutions in dimethyl sulfoxide (DMSO) were applied to the filter paper (20 µl per dish). 5 µl of *R. solanacearum* (OD_600_ = 0.5) was inoculated onto the NB medium, and plates were incubated at 30 ℃ for 48 h. After incubation, 1 ml of sterile water was used to wash the bacterial colonies, and then 50 µl of the bacterial suspension was diluted to 200 µl for OD_600_ measurements using a SpectraMax M5 spectrophotometer (Molecular Devices, Sunnyvale, CA, USA). DMSO was used as a blank control. All treatments were performed with three replicates.

### Data analysis and statistics

All data analyses were performed using R version 4.3.1. Microbiome community diversity was calculated as ASVs. Changes in community composition were assessed with a principal component analysis (PCA) by using the R package “microceo” v1.9.0 (54), function adonis in R package “vegan” v2.6.4 (55) was conducted to evaluate the significant difference in community structure across the different treatments. The “ONE_Tukey_HSD2” function was uesd to compare the differences between different treatments. The “LorMe” v1.1.0 package (56) in the laboratory was employed to construct microbial co-occurrence networks. The “ComplexHeatmap” v1.10.2 package (57) was utilized to depict the heatmap of bacterial resource utilization ability.

## Results

### Effect of inulin on promoting pathogen suppression of *L. sphaericus* HR92

To identify potential specific resource, we screened 13 water-soluble sugars based on the criterion that they should not directly promote pathogen growth while supporting the proliferation and antagonistic activity of beneficial bacteria. Initial in vitro assays revealed that, except for allulose, none of the tested water-soluble sugars significantly affected the growth of *R. solanacearum* (F₁₃,₂₈ = 8.48, p < 0.001; Fig. S2), indicating minimal risk of inadvertently stimulating pathogen development. We next assessed the effects of these 12 sugars on the beneficial strain *L. sphaericus* HR92 and its ability to inhibit *R. solanacearum*. Most sugars enhanced strain HR92 growth and strengthened its antagonistic activity, with inulin exhibiting the most pronounced effect. Specifically, inulin significantly promoted HR92 growth (F₈,₁₈ = 11.60, p < 0.001; Fig. S3A), with a maximum increase of 14.8%, and markedly improved its inhibitory effect against *R. solanacearum*, achieving the highest inhibition rate of 27.8% (F₈,₁₈ = 10.74, p < 0.001; Fig. S3B). Given its dual benefits, inulin was selected for subsequent experiments (Fig. S1).

In order to further verify effect of inulin on enhancing pathogen suppression of strain HR92, we conducted a greenhouse experiment. Inoculation of strain HR92 significantly reduce disease index of tomato plants, and inulin addition strengthened the disease suppression of strain HR92 with a dose effect (*F_3,12_* = 45.34, *p* < 0.05, Fig. 1A). The lowest disease index was observed in the treatment with 0.3% inulin and strain HR92, which reduced by a 70% reduction compared to the control (CK, Fig. 1B). Similarly, the *R. solanacearum* density in the rhizosphere of treatment with 0.3% inulin and strain HR92 was the lowest (*F_3,12_* = 46.95, *p* < 0.05, Fig. 1B). Interestingly, though 0.1% inulin increased the abundance of strain HR92 in the rhizosphere, higher inulin concentration inulin (0.3%) did not further bring strain HR92 to a higher density (*F_3,12_* = 13.223, *p* < 0.05, Fig. 1C). Additionally, we tested effect of inulin concentration on the growth of strain HR92. Consistent with the in planta findings, 0.1% inulin significantly increased the growth of strain HR92, while 0.3% showed no significant improvement (*F_3,8_* = 14.51, *p* < 0.05, Fig. S4). These findings indicate the effect of inulin on improving pathogen suppression of strain HR92, but not in a dose dependent way.

**Fig. 1.**
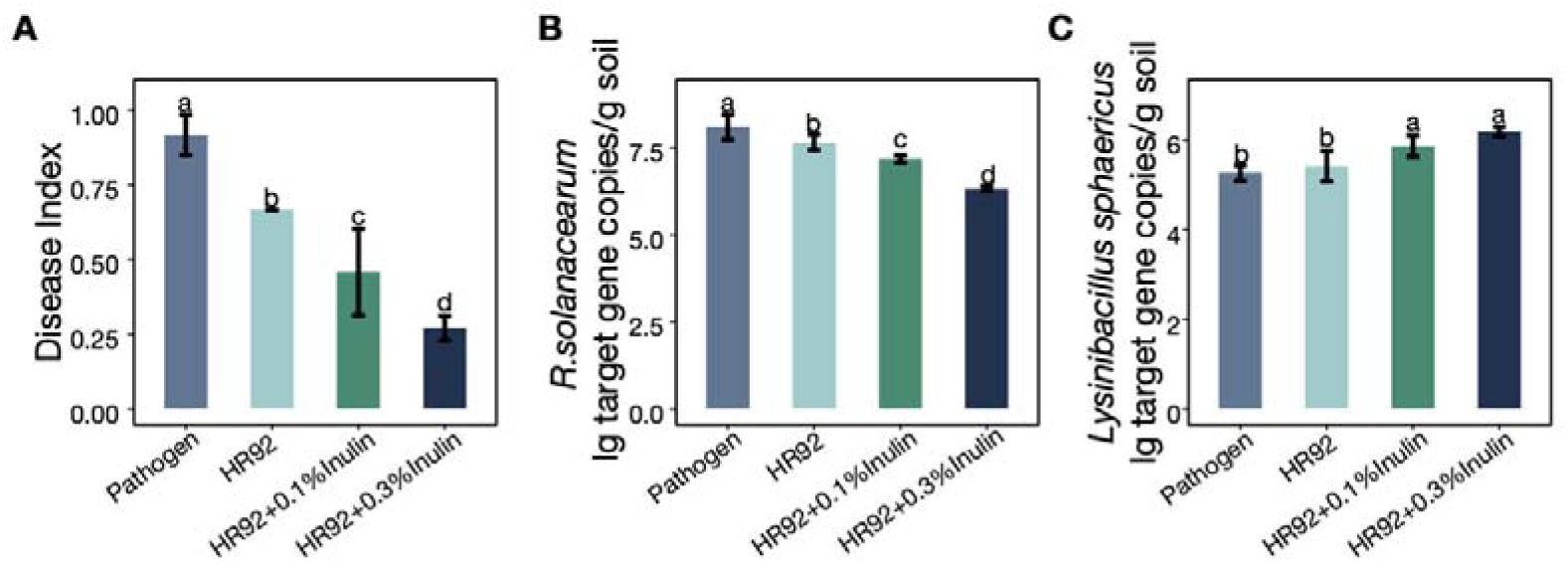
Effects of inulin on enhancing pathogen suppression by strain HR92 in tomato bacterial wilt. (A) Tomato disease index under different treatments (*N* = 4). (B) Absolute abundance of *R. solanacearum* in the tomato rhizosphere under different treatments, quantified by qPCR using species-specific primers (one-way ANOVA with correction by Tukey’s HSD test; *N* = 4). (C) Absolute abundance of HR92 in the tomato rhizosphere under different treatments, quantified by qPCR using species-specific primers (one-way ANOVA with correction by Tukey’s HSD test; *N* = 4). Different letters in the bar graph indicate significant differences (*p* < 0.05).

### Strain HR92 and inulin alter the bacterial community of the tomato rhizosphere

16S rRNA gene amplicon sequencing was conducted to reveal the differences in the rhizosphere bacterial community among the four treatments. There were no significant differences in the Simpson and Shannon diversity indices between the control group and the HR92 or HR92+0.1% inulin treatment groups. However, the addition of 0.3% inulin with strain HR92 significantly decreased both the Simpson and Shannon indices of the tomato rhizosphere bacterial community (Shannon: *F_3,12_* = 9.26, *p* = 0.02, Fig 2A; Simpson *F_3,12_* = 3.36, *p* = 0.04, Fig 2B).

**Fig. 2.**
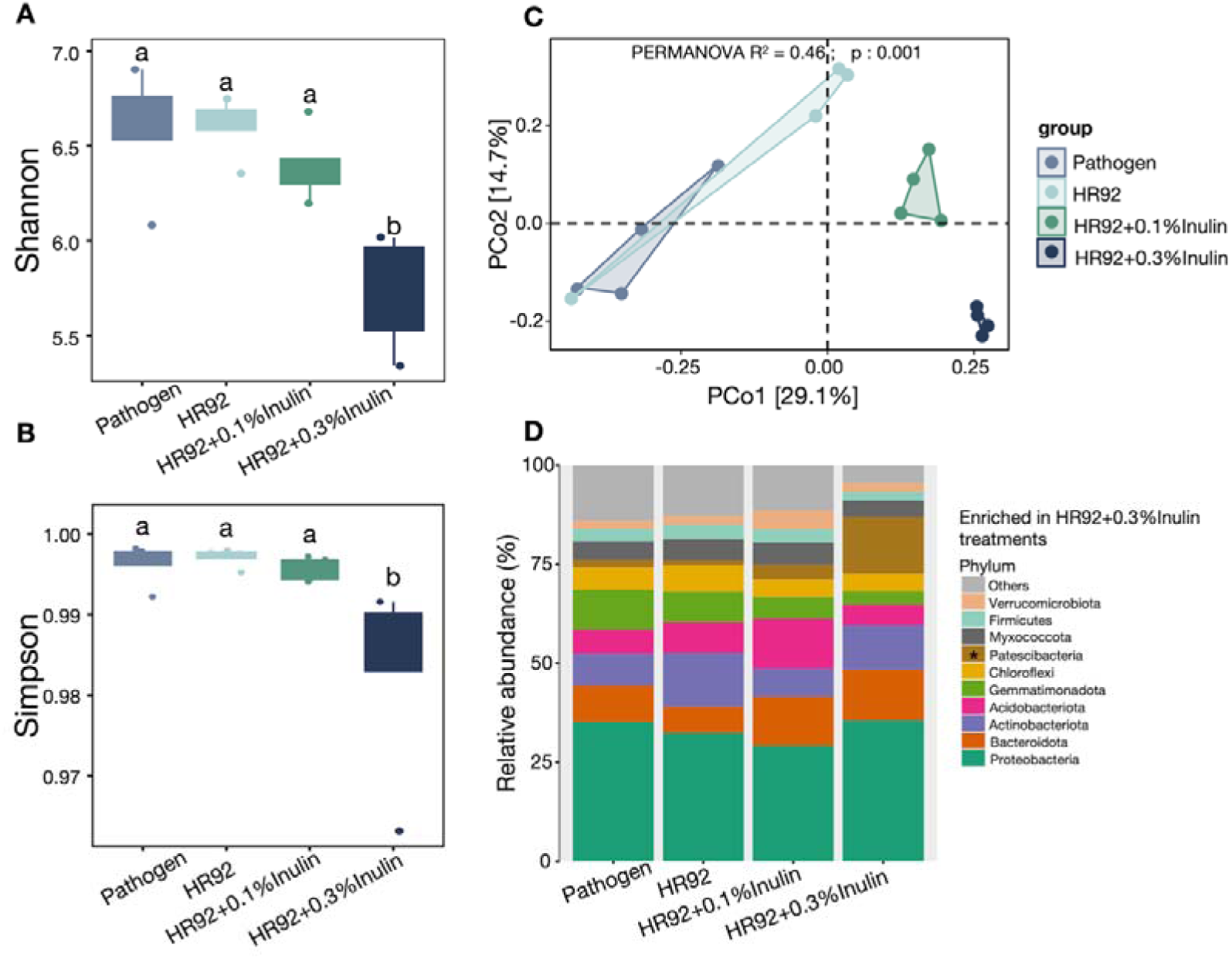
Microbial community analysis under different treatments. (A) Alpha diversity analysis of the rhizosphere microbiome across treatments using the Shannon index. (B) Alpha diversity analysis using the Simpson index. (C) PCoA of the rhizosphere soil microbiota under each treatment. (D) Relative abundance of the top 10 most abundant phyla in the rhizosphere soil across treatments. The asterisk indicates phyla enriched specifically in the HR92+0.3% inulin treatment compared to the other treatments. Different letters above the boxes of each compartment indicate a significant difference at *p* < 0.05 (one-way ANOVA with correction by Tukey’s HSD test).

Principal coordinates analysis (PCoA) revealed significant differences in the bacterial community composition among the four treatments. Inoculating HR92 can alter the tomato rhizosphere microbial community, and the addition of inulin can further modify the microbial community structure, with distinct differences observed between the 0.1% and 0.3% inulin treatments, being especially clear along the PCo1, which explained 29.1% of the total variation in bacterial community composition (PERMANOVA, *R^2^* = 0.460, *p* = 0.001, Fig. 2C).

Further analysis of the relative abundance of bacteria in different treatments revealed that bacteria amplicon sequence variants (ASVs) primarily belonged to 10 phyla, including *Proteobacteria*, *Acidobacteria*, *Bacteroidetes*, *Verrucomicrobia*, *Chloroflexi*, *Actinobacteria*, *Patescibacteria*, *Gemmatimonadetes*, *Myxococcota*, and *Firmicutes*. Although the composition of dominant bacterial phyla was similar across treatments, the relative abundance of certain dominant phyla differed significantly. Specifically, the phylum *Patescibacteria* was significantly enriched in the treatment with 0.3% inulin and strain HR92 (Fig. 2D).

### Inulin stimulates native soil resident microbes to assist strain HR92 in resisting *R. solanacearum* invasion

Following the application of 0.3% inulin in combination with HR92, the colonization of HR92 in the rhizosphere did not significantly increase (Fig. 1C). However, the disease index of tomatoes and the population of rhizosphere *R. solanacearum* further decreased (Fig. 1A, B). We hypothesize that, beyond its direct effect on HR92, the 0.3% inulin may enrich certain native soil resident microbes that contribute to pathogen suppression. Subsequently, 21 genera significantly enriched in the treatment with 0.3% inulin were identified, defining them as potential key taxa (Table S1, *p* < 0.05).

Further analysis of co-occurrence networks and distribution patterns of dominant genera across the four treatments identified key taxa. Notably, among the top ten nodes in the co-occurrence network based on betweenness centrality, two ASVs, ASV2 (*Pseudarthrobacter*) and ASV201 (*Agromyces*), were recognized as potential keystone bacteria (Fig. 3A). The relative abundance of ASV2 was significantly correlated with inulin concentration, with its abundance further increasing under the 0.3% inulin treatment, whereas ASV201 was enriched exclusively in the 0.3% inulin treatment (ASV2: *F_3,12_* = 13.16, *p* < 0.001; ASV201: *F_3,12_* = 19.18, *p* < 0.001, Fig. 3B). Both ASV2 (R = −0.82, *p* < 0.05) and ASV201 (R = −0.81, *p* < 0.05) exhibited significant negative correlations with the abundance of *R. solanacearum*, and *Pseudarthrobacter* (*R* = −0.77, *p* < 0.05) and *Agromyces* (*R* = −0.75, *p* < 0.05) also demonstrated significant negative correlations with the abundance of *R. solanacearum* (Fig. S5A, B). Thus, *Pseudarthrobacter* and *Agromyces* are considered potential key taxa that interact synergistically with HR92 to confer disease resistance. Additionally, microorganisms enriched under the 0.3% inulin treatment were predominantly concentrated within modules 2 and 3 of the co-occurrence network, both of which were significantly negatively correlated with pathogen abundance (module 2: *R* = −0.69, *p* < 0.05; module 3: *R* = −0.82, *p* < 0.05; Fig. S5C, D), suggesting their crucial roles in disease suppression.

**Fig. 3.**
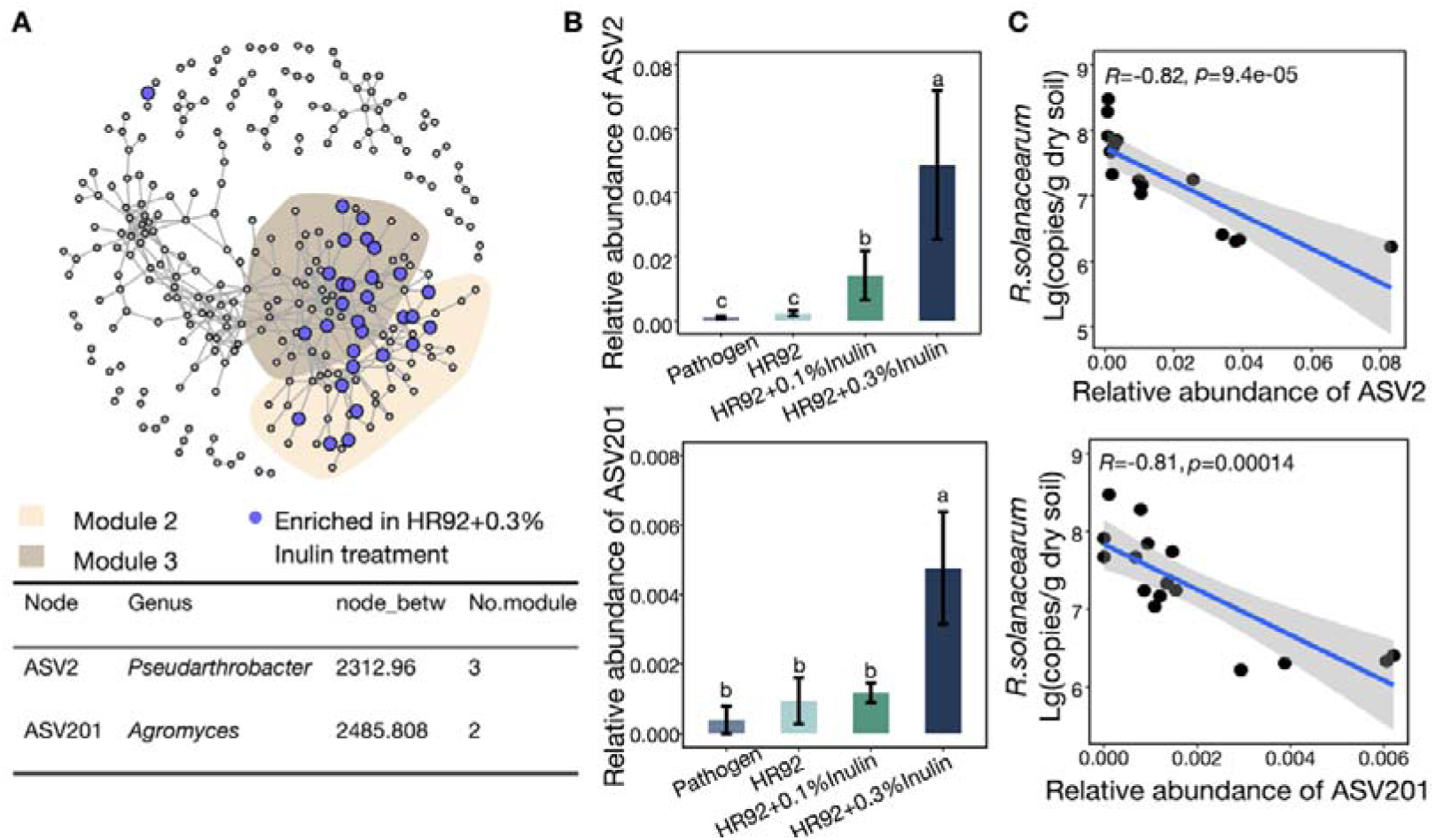
Co-occurrence network analysis. (A) The co-occurrence network at the ASV level, where the purple nodes represent ASVs enriched under high-concentration inulin treatment. Key nodes and their properties are detailed below the network graph. (B) Relative abundance of ASV2 and ASV201 in the rhizosphere microbial communities across four treatments (*N* = 4; one-way ANOVA with correction by Tukey’s HSD test). (C) Linear regression between the relative abundance of ASV2 and ASV201 and the population of *R.solanacearum* in the tomato rhizosphere (*p* < 0.05). Different letters in the graph indicate significant differences (*p* < 0.05).

Tax4Fun functional prediction indicated that *Pseudarthrobacter* is primarily associated with secondary metabolite synthesis, basic metabolism, and isoflavonoid biosynthesis, suggesting that *Pseudarthrobacter* may mediate disease resistance via these pathways (Fig. S6). Given its significant association with disease suppression and its tenfold higher abundance compared to *Agromyces* in the tomato rhizosphere, *Pseudarthrobacter* likely plays a more prominent role in mediating rhizosphere interactions. Therefore, subsequent validation experiments focused on *Pseudarthrobacter*.

### *Pseudarthrobacter* strains in cooperative interaction with HR92 to suppress *R. solanacearum* invasion

To evaluate the effect of *Pseudarthrobacter* on disease suppression and explore its underlying mechanisms, we isolated 50 culturable bacterial strains from soil treated with 0.3% inulin. Among these, strains P265 and P397 were identified as members of the genus *Pseudarthrobacter*. These two strains were used to investigate the specific mechanisms involved. First, we examined the effect of inulin on the growth of the two bacterial strains and found that inulin significantly promoted the growth of strain P265 and P397 (strain P265: t-test: *t_6.32_* = −2.82, *p* = 0.029; strain P397: t-test: *t_8.67_* = −6.42, *p* < 0.001; Fig. 4A), indicating that these bacteria effectively utilized inulin as a resource. We subsequently assessed the potential of strain P265 and P397 to inhibit pathogenic activity. In vitro assays demonstrated that the supernatants from P265 and P397 did not directly inhibit *R. solanacearum* (strain P265: *F_2,11_* = 0.69, *p* = 0.5; strain P397: *F_2,11_* = 1.15, *p* = 0.5, Fig. S8). However, under co-culture conditions, both P265 and P397 inhibited the growth of *R. solanacearum*, the inhibition rate of P265 was 41%, and that of P397 was 41.4%, with the addition of inulin showing no significant influence on this inhibitory effect (strain P265: *F_2,9_* = 130.13, *p* < 0.001; strain P397: *F_2,9_* = 56.58, *p* < 0.001; Fig. 4B). These observations suggest that the inhibitory mechanism may involve nutrient competition between these strains and *R. solanacearum*.

**Fig. 4.**
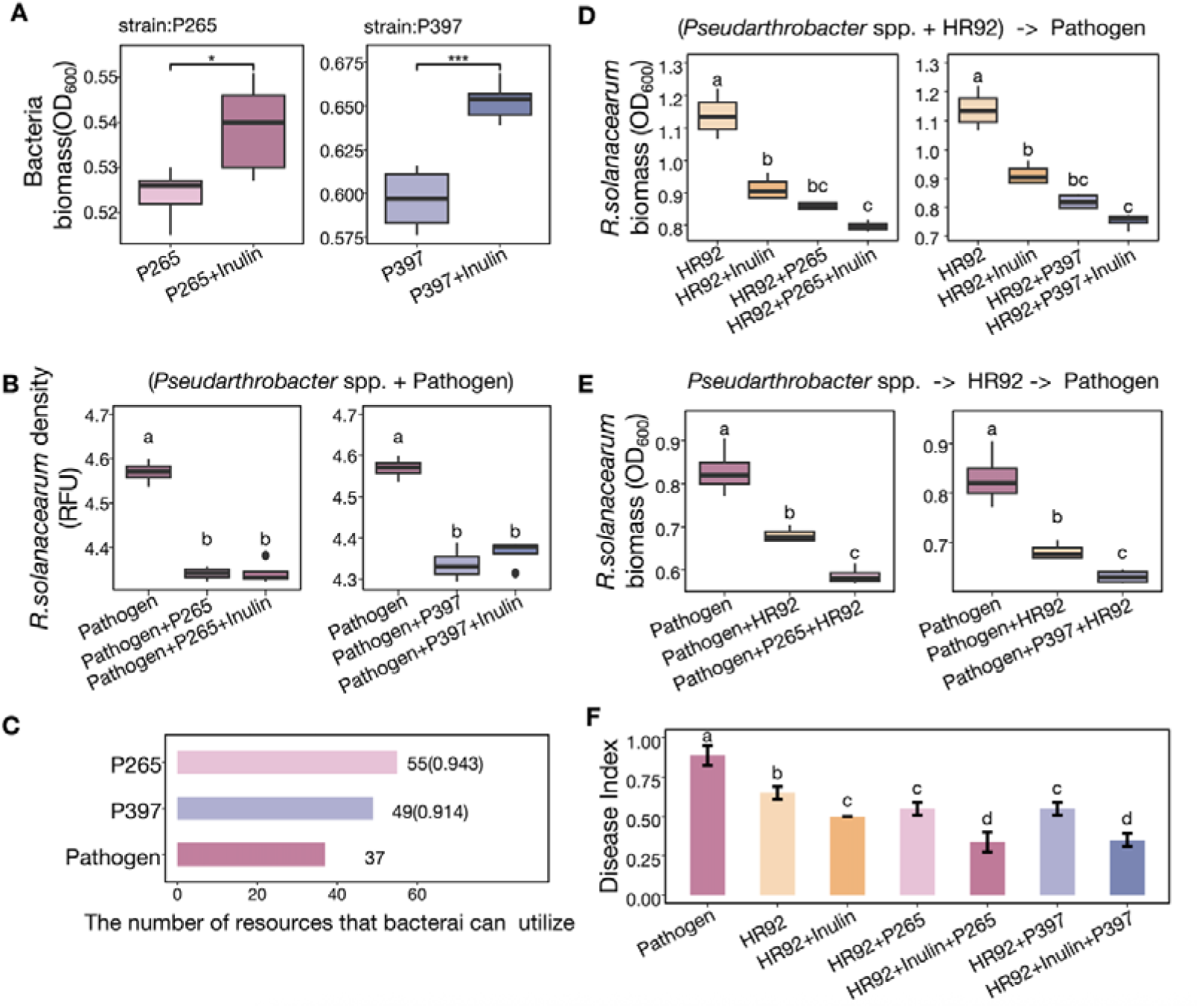
Investigation of the antibacterial effects of *Pseudarthrobacter* spp. and its influence on the antibacterial activity of HR92. (A) Effect of inulin on the growth of *Pseudarthrobacter siccitolerans* (P265) and *Pseudarthrobacter oxydans* (P397). Asterisk means *p* < 0.05 and three asterisks means *p* < 0.001 under student’s t test (*N* = 4). (B) Inhibition of *R. solanacearum* by strains P265 and P397 under co-culture conditions with and without the addition of inulin. (C) Resource utilization overlap between strain P265/P397 and *R. solanacearum*, indicating the potential nutritional competition. (D) Enhanced antibacterial activity of the supernatant after co-culture of strain HR92 with strains P265 and P397. The arrows at the top of the box plot represent the feeding procedure. Specifically, they indicate feeding the pathogen with the supernatant from co-cultures of strain P267 or P397 and strain HR92. (E) Increased antibacterial activity of HR92 after being cultured with the supernatants from strains P265/P397. The arrows at the top of the box plot represent the feeding procedure. Specifically, they indicate feeding strain HR92 with the supernatant from strain P267 or P397, followed by feeding the pathogen with the supernatant from strain HR92. (F) Pot experiments showing enhanced tomato disease resistance when HR92, *Pseudarthrobacter* strains, and inulin were applied together. Different letters in the graph indicate significant differences (*p* < 0.05).

To explore nutrient competition we examined the utilization of 74 different rhizosphere carbon sources by P265, P397, and *R. solanacearum*. The results indicated that P265 and P397 utilized a broader range of resources compared to *R. solanacearum* (Table 2, *p* < 0.05). The overlap in resource utilization between P265 and *R. solanacearum* was 0.94, and between P397 and *R. solanacearum* was 0.91 (Fig. 4C), indicating potential competition for carbon resources between *Pseudarthrobacter* strains and *R. solanacearum*.

Additionally, the supernatant from the co-culture of strain HR92 with P265 or P397 showed an enhanced inhibitory effect against *R. solanacearum* (strain P265: *F_3,12_* = 19.18, *p* < 0.001; strain P397: *F_3,12_* = 22.35, *p* < 0.001; Fig. 4D). We hypothesized that either strain HR92 induced antibacterial activity in strains P265 and P397 or that these strains augmented strain HR92’s antibacterial function. To test this hypothesis, first, we fed the supernatant of strain HR92 to P265 and P397. Subsequently, we fed the supernatants of P265 and P397 to *R. solanacearum* and found that the growth of *R. solanacearum* was not affected (strain P265: *F_2,18_* = 1.834, *p* = 0.18; strain P397: *F_2,18_* = 1.55, *p* = 0.24, Fig. S9). This indicates that strain HR92 was unable to confer the ability to antagonize *R. solanacearum* to P265 or P397. Conversely, we fed the supernatant of strain P265 or P397 to strain HR92 and found that the supernatant of strain HR92 had a stronger ability to inhibit *R. solanacearum*, with enhancements of 12% and 6.6%, respectively (strain P265: *F_2,9_* = 46.23, *p* < 0.001; strain P397: *F_2,9_* = 33.306, *p* < 0.001; Fig. 4E). In pot experiments, consistent with the in vitro findings, strains P265 and P397 enhanced the ability of strain HR92 to mitigate tomato bacterial wilt. Notably, the combined treatment of strain HR92, *Pseudarthrobacter* strains, and inulin provided the best protection against tomato bacterial wilt, with a disease index of 0.3375 (P265) and 0.35 (P397), achieving control efficiencies of 61.9% (P265) and 60.5% (P397) (*F_6,21_* = 87.21, *p* < 0.001, Fig. 4F).

### *Pseudarthrobacter* strains and inulin enhance cyclic lipopeptide surfactin and 3-methyl-1-butanol production by HR92

Transcriptomic analysis was conducted to elucidate the individual and combined effects of inulin and *Pseudarthrobacter* strains on strain HR92. Inulin upregulated genes involved in flagellar assembly, bacterial chemotaxis, fatty acid biosynthesis, and siderophore biosynthesis in strain HR92, indicating that inulin enhances the motility of HR92 in the rhizosphere, thereby increasing its colonization quantity(Fig. S10). Strains P265 and P397 collectively influenced genes expression in strain HR92, with significant differential expression observed in genes related to the synthesis of 4-aminobutyrate, methyl-accepting, butanediol and bacitracin, all of which are associated with strain HR92’s antimicrobial activity (Fig. S11). Bacitracin is a broad-spectrum antibiotic known for its antimicrobial properties. Furthermore, the combined action of *Pseudarthrobacter* strains and inulin enhanced the synthesis of amino acids such as valine and leucine in HR92 (Fig. S12, Fig. S13). Subsequent Weighted Gene Co-expression Network Analysis (WGCNA) identified three gene modules in strain HR92 that were linked to its antimicrobial activity (Fig. S14). Within the black and pink modules, two genes highly correlated with the modules and exhibiting the highest correlation with strain HR92’s antimicrobial effect were identified: LYS2002816 and LYS2001501. These genes were associated with butanediol and NAD(P)H, both of which exhibited a significant positive correlation with strain HR92’s antimicrobial capability (LYS2002816: *R* = 0.79, *p* <0.05, LYS2001501: *R* = 0.73, *p* <0.05, Fig. S15). The abundance of these genes was highest in strain P265 or P397, HR92, and inulin treatment (LYS2002816: *F_5,12_* = 2.9, *p* < 0.05, Figure 5A; LYS2001501: *F_5,12_* = 5.8, *p* < 0.05, Figure 5B). Literature review indicated that butanediol contributes to the synthesis of the volatile antimicrobial compound 3-methyl-1-butanol in strain HR92, whereas fatty acids, NAD(P)H, valine, and leucine are involved in the synthesis of the cyclic lipopeptide surfactin (Fig. 5C, D), ultimately enhancing strain HR92’s ability to inhibit *R. solanacearum*.

**Fig. 5.**
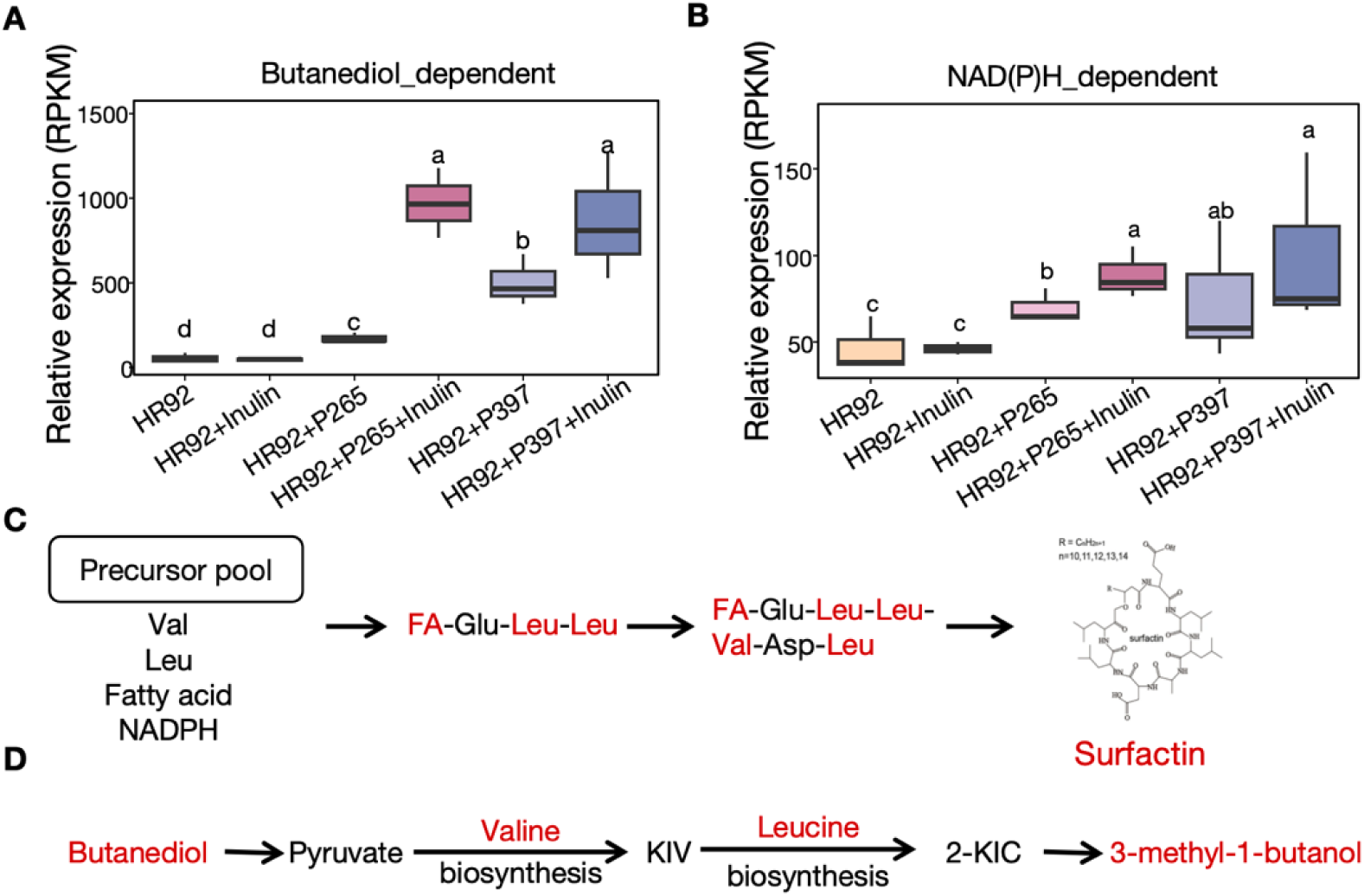
Synergistic enhancement of HR92 antibacterial capability by *Pseudarthrobacter* spp. and inulin. (A) Expression levels of HR92 genes associated with butanediol synthesis across six treatments, quantified by RPKM tables. (B) Expression levels of HR92 genes involved in NAD(P)H synthesis across six treatments, quantified by RPKM tables. Different letters in the graph indicate significant differences (Duncan’s multiple range test). (C) Schematic diagram illustrating the involvement of fatty acids, NAD(P)H, valine, and leucine in the biosynthesis of the cyclic lipopeptide surfactin. (D) Schematic diagram showing the pathway through which butanediol is converted to 3-methyl-1-butanol via pyruvate, valine biosynthesis, and leucine biosynthesis pathways. Red font highlights the pathways or genes influenced by the synergistic action of *Pseudarthrobacter* spp. and inulin.

To further elucidate the role of 3-methyl-1-butanol, we tested its effect on *R. solanacearum* growth. Our results showed that 50% and 100% concentrations of 3-methyl-1-butanol could inhibit the growth of *R. solanacearum*, with 50% concentration resulting in a 16.4% inhibition rate and 100% concentration achieving a 40% inhibition rate (*F_4,10_* = 6.839, *p* < 0.05, Fig. S16).

## Discussion

There is growing interest in harnessing beneficial bacteria for plant disease biocontrol, yet strategies to maxmize their effectiveness remain insufficiently explored. Here, our study emphasizes the important role of inulin in promoting the growth and pathogen-suppressing capabilities of the beneficial strain HR92, along with its synergistic interactions with native soil microbes to manage *R. solanacearum* infections in tomatoes. We also investigated whether increased inulin concentrations could further improve the biocontrol efficacy of strain HR92. Both *in vitro* and *in vivo* experiments demonstrated that inulin positively influenced bacterial colonization and biocontrol efficiency. In general, higher inulin concentrations selectively enriched *Pseudarthrobacter* species in the soil, which acted synergistically with strain HR92 to enhance biocontrol outcomes.

Inulin promoted strain HR92 growth and augmented its pathogen-suppressing ability. Initially, we observed a threefold increase in strain HR92 population in the tomato rhizosphere upon inulin treatment. This finding is consistent with previous studies showing that inulin selectively stimulates the growth of beneficial bacteria, such as *Bifidobacterium*, *Lactobacillus*, and *Bacteroides,* while inhibiting potential pathogens (58). Additionally, inulin has been shown to increase the production of lactic acid, acetic acid, and other volatile compounds by probiotic strains, indicating a positive synergistic interaction between inulin and beneficial microbes (59). These findings support our results that inulin, as a prebiotic, promotes the growth of beneficial bacteria. We further discovered that the disease severity index for tomatoes treated with both inulin and strain HR92 decreased by 31.2% compared to those treated with strain HR92 alone. Previous research has demonstrated that strain HR92 could effectively control *R. solanacearum* (19), suggesting a direct impact of strain HR92 on the *R. solanacearum*. This can be interpreted as the biocontrol efficacy of beneficial bacteria being closely related to their abundance in the rhizosphere (60), and that plant-derived carbon sources could enhance the metabolic activity of specific beneficial bacteria, thereby improving their disease resistance (61, 62). Furthermore, we observed that increasing the inulin concentration from 0.1% to 0.3% did not further promote the growth of strain HR92, which may be due to strain HR92 already obtaining sufficient nutrients at the 0.1% concentration. The optimal inulin concentration for promoting strain HR92 growth under complex environmental conditions remains to be further investigated.

Inulin and its varying concentrations significantly influenced rhizosphere microbial diversity and community structure. While a low inulin concentration (0.1%) had no significant effect on the Simpson and Shannon diversity indices, a higher concentration (0.3%) reduced these indices. This can be explained by higher concentrations of inulin potentially altering community structure by selectively enriching certain functional bacteria, thereby reducing the relative abundance of other bacterial groups. Our findings revealed distinct differences in bacterial community composition between the two inulin concentrations treatments. Previous studies have indicated that the concentration of exogenous carbon sources in soil has a significant selective effect on native microbial communities (40, 63). Taken together, different concentrations of inulin can regulate the interactions and structure of native microbial communities, maintaining higher diversity at lower concentrations while promoting the enrichment of specific functional bacteria at higher concentrations. Since a higher concentration of inulin further reduced disease severity and the abundance of *R. solanacearum* in the rhizosphere, without altering the abundance of strain HR92 in the rhizosphere. Therefore, we hypothesize that, beyond directly benefiting strain HR92, a 0.3% inulin concentration may also enrich certain native soil microbes that aid in suppressing pathogens. In fact, in the co-occurrence network analysis, *Pseudarthrobacter* and *Agromyces* were significantly enriched under high inulin concentration treatment and identified as key network hubs, rowten showing a significant negative correlation with the pathogen. This suggests that they may support strain HR92’s pathogen suppression function through competition or antagonistic interactions. Interestingly, previous studies have indicated that *Pseudarthrobacter* is adept at collaborating with other beneficial bacteria to perform better ecological functions (64). On the other hand, strains belonging to the genus *Pseudarthrobacter* are known as copiotrophic root colonizers (65, 66) and have been proposed as a key members of the root microbiome (67). In summary, the present study highlights the importance of the synergistic interaction between inulin, *Pseudarthrobacter* and HR92 in managing *Ralstonia solanacearum*.

Our study also demonstrates that the combined effects of inulin and *Pseudarthrobacter* strains significantly enhance the antibacterial activity of HR92 against *R. solanacearum*, primarily through the upregulation of genes related to the synthesis of key metabolites, such as surfactin and 3-methyl-1-butanol. The inhibitory effects of these metabolites on pathogens have been well-documented. The cyclic lipopeptide surfactin, a polypeptide compound composed of fatty acids and amino acids, exhibits notable antimicrobial, antiviral, and plant stress-resistance-promoting activities, playing a particularly prominent role in disease defense (68). Transcriptomic analysis in this study revealed that inulin and *Pseudarthrobacter* strains promoted the synthesis of surfactin by upregulating metabolic pathways related to fatty acids and amino acids, such as valine and leucine. Furthermore, 3-methyl-1-butanol, as a volatile antimicrobial compound, has shown high efficacy in inhibiting various pathogens. Its mode of action involves binding to microbial cell membranes, inserting into the bilayer structure, and altering membrane permeability, which destabilizes the membrane, causes leakage of cellular contents such as ions and ATP, and ultimately leads to cell death, effectively inhibiting pathogen growth (69, 70). Our co-expression network analysis of HR92 genes indicated that the butanediol and NADPH metabolic pathways are closely associated with the antibacterial potential of HR92, possibly by promoting the synthesis of 3-methyl-1-butanol. Further experimental validation confirmed that high concentrations of 3-methyl-1-butanol exhibit strong inhibitory effects on *R. solanacearum*, which is consistent with other studies on volatile organic compounds (VOCs) as disease suppressants (71). In summary, these results suggest that the synergistic effects of inulin and *Pseudarthrobacter* can significantly enhance HR92’s ability to suppress *R. solanacearum* by promoting the synthesis of key antibacterial metabolites.

In conclusion, our study demonstrates that the inulin significantly enhances strain HR92 colonization, chemotaxis, fatty acid metabolism, and siderophore production in tomato roots. The addition of 0.3% inulin had a greater impact than 0.1%, notably enriching *Pseudarthrobacter* species within the rhizosphere. These species promoted the synthesis of butanediol and bacitracin in strain HR92, thereby enhancing its antimicrobial capabilities. Furthermore, *Pseudarthrobacter* spp. competed with *R. solanacearum* for root-associated resources, reducing the abundance of pathogenic bacteria via nutrient competition. The combined effects of *Pseudarthrobacter* spp. and inulin further upregulated strain HR92’s valine and leucine metabolic pathways. Valine, leucine, fatty acids, and NAD(P)H are essential precursors for *L. sphaericus* to synthesize the antimicrobial cyclic lipopeptide surfactin. Additionally, butanediol is converted into the antimicrobial VOC 3-methyl-1-butanol through valine and leucine metabolic pathways. Thus, the cooperative interaction between *Pseudarthrobacter* spp. and inulin enhanced strain HR92’s ability to inhibit *R. solanacearum*, ultimately reducing the incidence of tomato disease. This research provides valuable insights into the synergistic interactions between exogenous additives and beneficial microbes, offering a foundation for developing advanced, targeted formulations for sustainable and effective disease management (Fig 6).

**Fig. 6.**
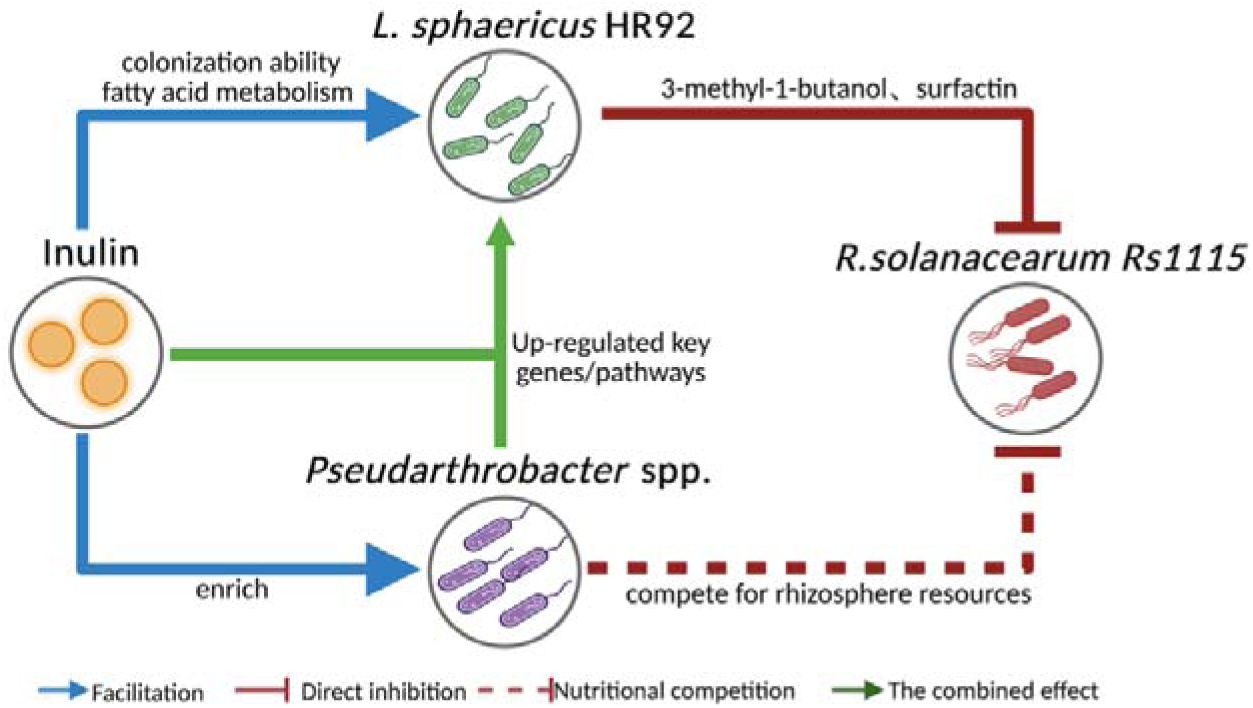
Pathways by which inulin enhances HR92’s disease-resistance mechanisms. Blue arrows indicate promotion effects, red solid lines indicate direct inhibition effects, red dashed lines indicate inhibition of pathogen growth through nutrient competition.

## Data availability

All metagenomics reads are publicly available in the SRA database under the accession number PRJNA1198698.

## Acknowledgments

This work was financially supported by the National Key Technology R&D Program of China (2023YFD1701505), the National Key Research and Development Program of China (2024YFC3406003), Guangxi Science and Technology Program (GuiKe AA24010003), the National Nature Science Foundation of China (42477125, 42090060, 42325704), the Natural Science Foundation of Jiangsu Province (BK20230988), the Technology Innovation Special Fund of Jiangsu Province for Carbon Dioxide Emission Peaking and Carbon Neutrality (BE2022423).

## Conflict of Interest

The authors declare no competing interests.

## Notes

### Competing Interest Statement

The authors have declared no competing interest.

### Summary of Updates

Revise the main figure and the corresponding description in the Results section

