## Supplemental fig1 for "Inulin reduces *Ralstonia* invasion by promoting cooperation between *Lysinibacillus sphaericus* and *Pseudarthrobacter* spp."

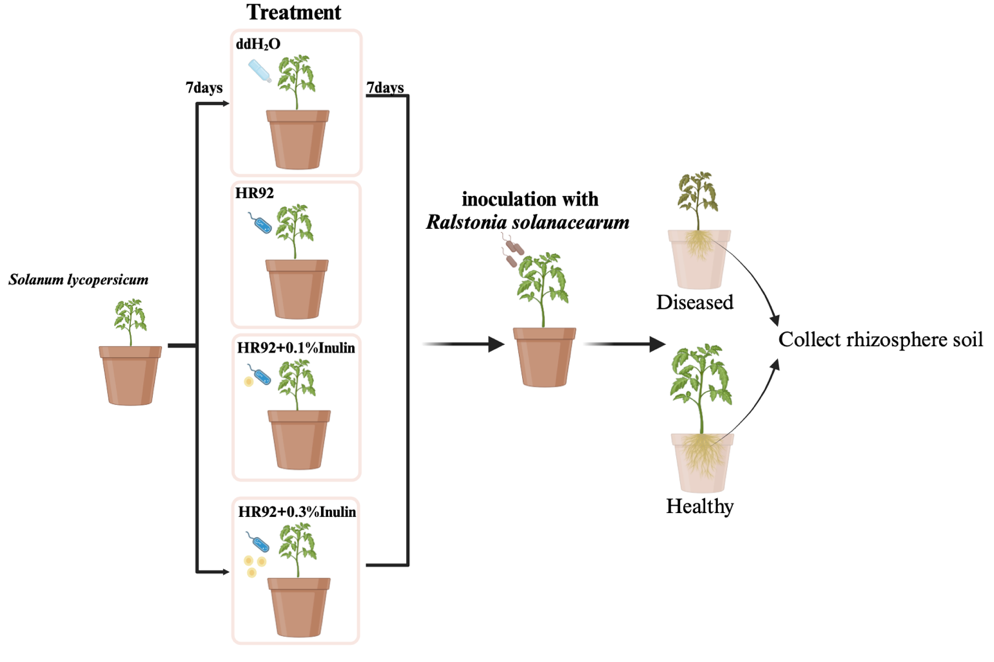

Fig. S1 Design of the pot experiment.

The potted plants were subjected to four treatments: (1) Only pathogen: Water was added seven days after transplanting tomatoes, followed by pathogen inoculation seven days later. (2) HR92: The beneficial bacterium HR92 was inoculated seven days after transplanting tomatoes, followed by pathogen inoculation seven days later. (3) HR92+0.1% Inulin: HR92 and inulin (1 g kg^-1^ soil) were added seven days after transplanting tomatoes, followed by pathogen inoculation seven days later. (4) HR92+0.3% Inulin: HR92 and inulin (3 g kg^-1^ soil) were added seven days after transplanting tomatoes, followed by pathogen inoculation seven days later.

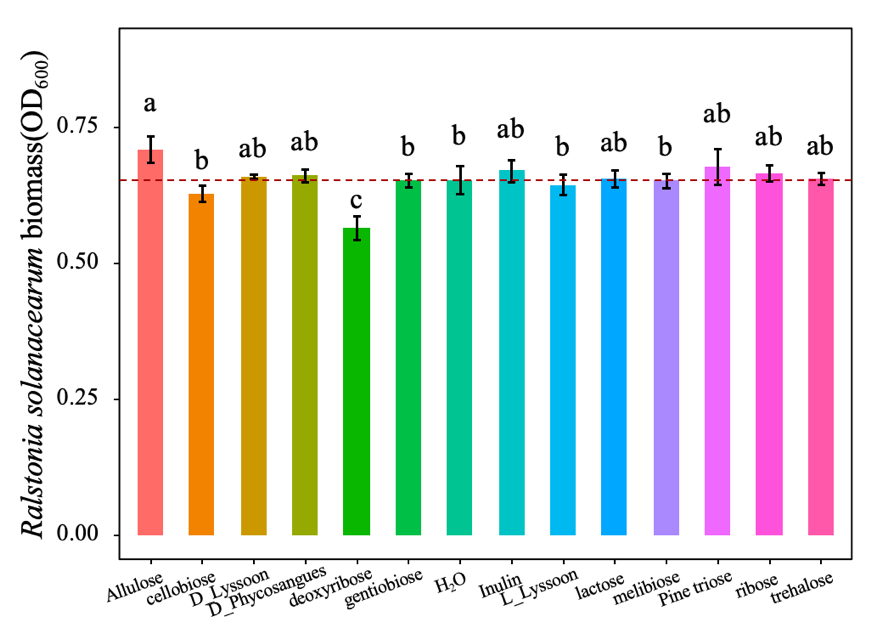

Fig. S2 The effect of different water-soluble sugars on the growth of *R. solanacearum.*

The red dashed line represents the growth of *R. solanacearum* without exogenous sugar addition. Different letters indicate significant differences (*F_13,28_* = 8.483, *p* < 0.001, *N* = 4).

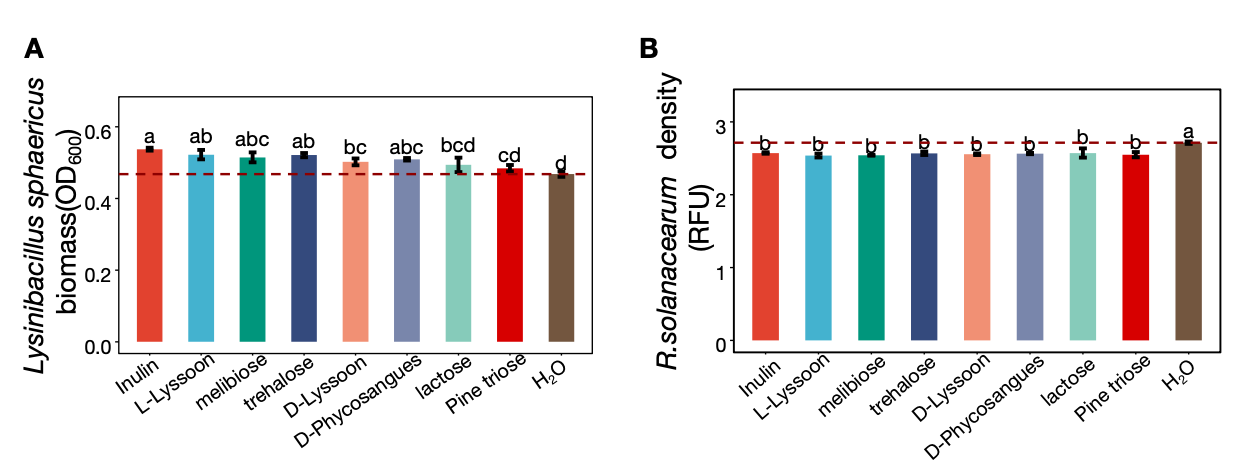

Fig. S3 The effect of different water-soluble sugars on the growth of HR92 (F₈,₁₈ = 11.60, p < 0.001) and its inhibitory activity against *R. solanacearum* (F₈,₁₈ = 10.74, p < 0.001). The red dashed line represents the growth of HR92 without the addition of water-soluble sugars (equal volume of sterile water added as control)

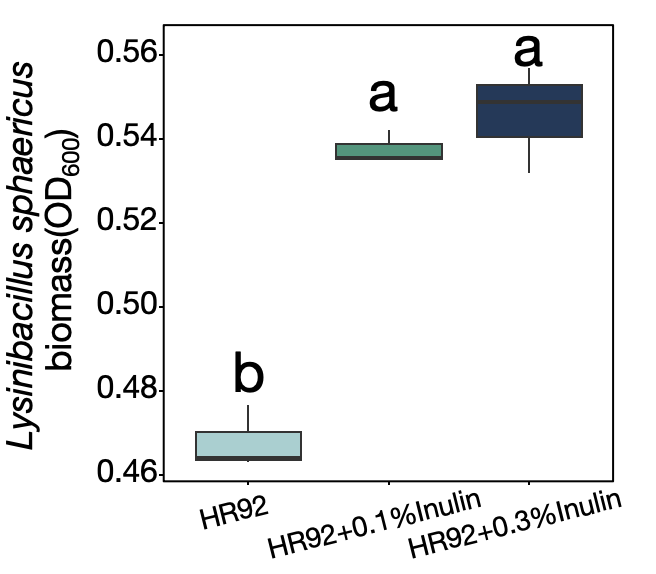

Fig. S4 Effect of different inulin concentrations on the growth of HR92. Different letters indicate significant differences (*F_3,8_* = 14.51, *p* < 0.05 , *N* = 4).

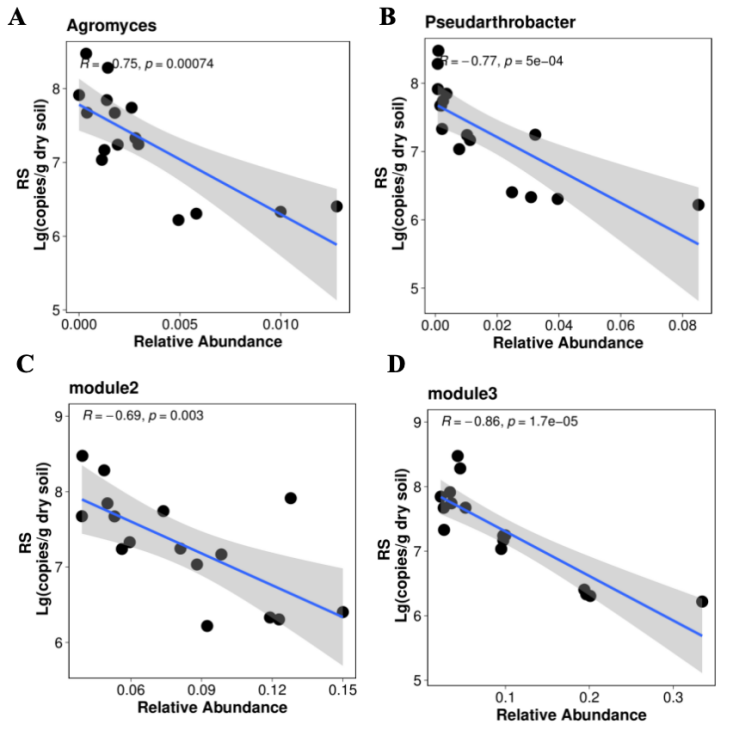

Fig. S5 Linear regression analysis between microbial relative abundance and R. solanacearum in tomato rhizosphere soil. (A) Linear regression between the relative abundance of *Agromyces* spp. and *R.solanacearum* in the rhizosphere soil of tomato. (B) Linear regression between the relative abundance of *Pseudarthrobacter* spp. and *R.solanacearum* in the rhizosphere soil of tomato. (C) Linear regression between the relative abundance of module2 and *R.solanacearum* in the rhizosphere soil of tomato. (D) Linear regression between the relative abundance of module3 and *R.solanacearum* in the rhizosphere soil of tomato. “R” stands for the correlation coefficient and “*p*” stands for the p-value

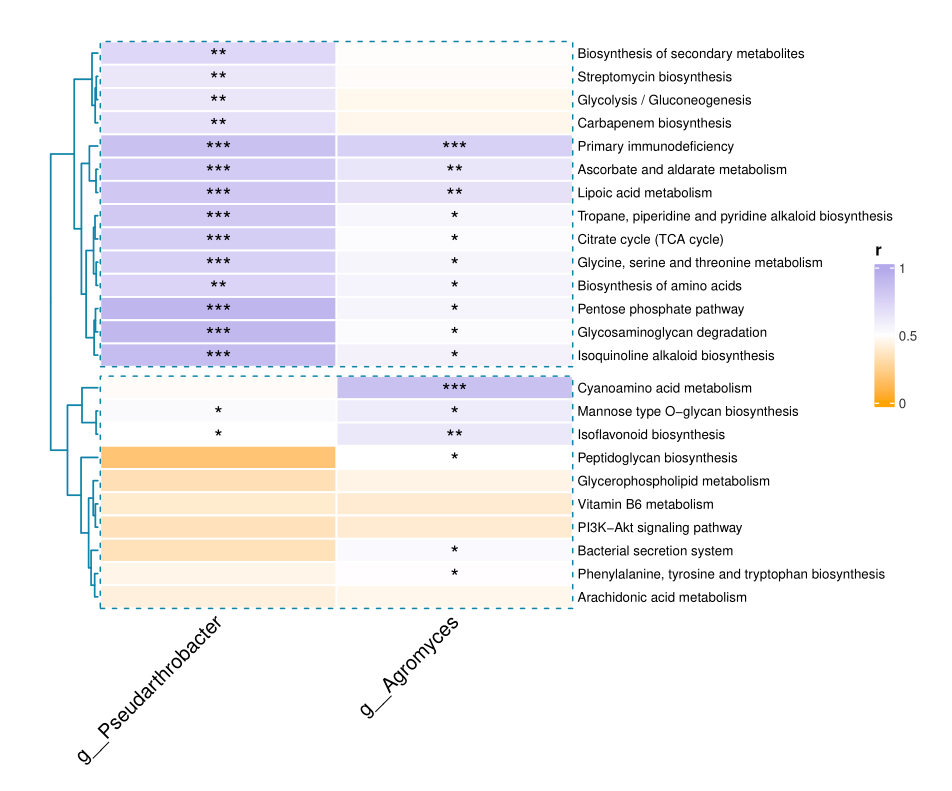

Fig. S6 Correlation between *Agromyces* spp. or *Pseudarthrobacter* spp. and community functions (Tax4Fun). The color of the heatmap cells represents the correlation coefficient. A sterisk means *p* < 0.05 , two asterisks means *p* < 0.01 and three asterisks means *p* < 0.001 under student’s t test.

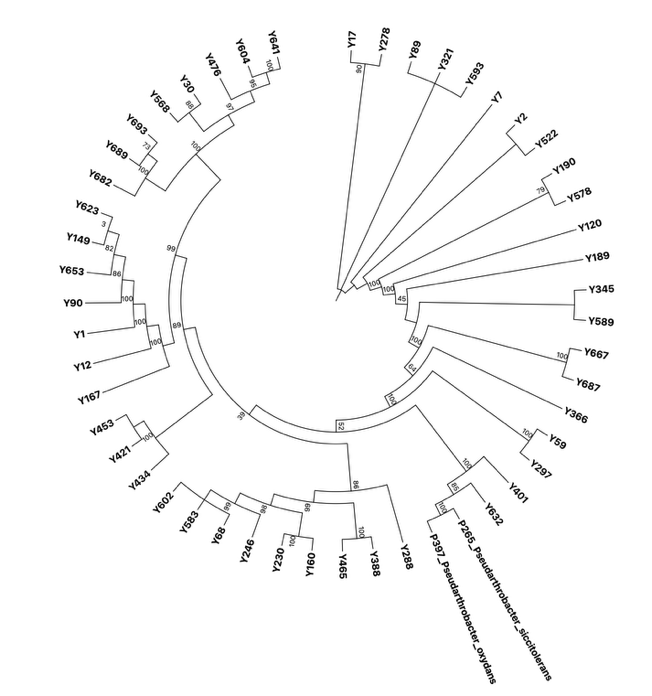

Fig. S7 Phylogenetic tree showing phylogenetic relationship among 50 isolated colonies. Full-length sequences of these isolates and representative sequences from 16S rRNA amplicons identified as *Pseudarthrobacter* spp. were aligned and used to construct a phylogenetic tree based on the maximum likelihood method. Numbers at the branches denote bootstrap values for the corresponding branches.

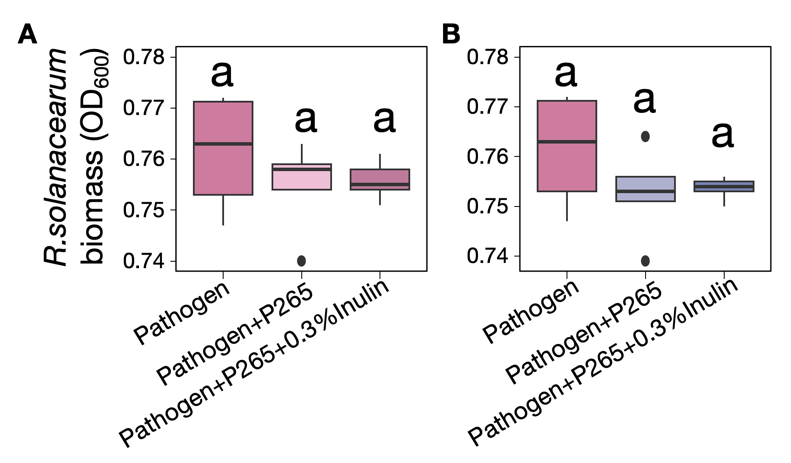

Fig. S8 Effects of Pseudarthrobacter supernatants on *Ralstonia* growth induced by inulin. Different letters indicate significant differences (strain P265: *F_2,11_* = 0.69, *p* = 0.5; strain P397: *F_2,11_* = 1.15, *p* = 0.5, *N* = 4).

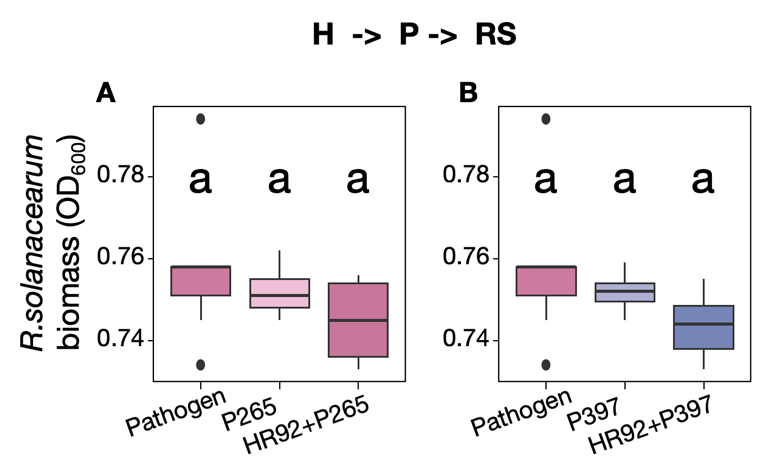

Fig. S9 Effect of HR92 on the inhibition activity of P265 and P397 against *R. solanacearum.* Different letters indicate significant differences (strain P265: *F**_2,18_* = 1.834, *p* = 0.18; strain P397: *F_2,18_* = 1.55, *p* = 0.24, *N* = 4).

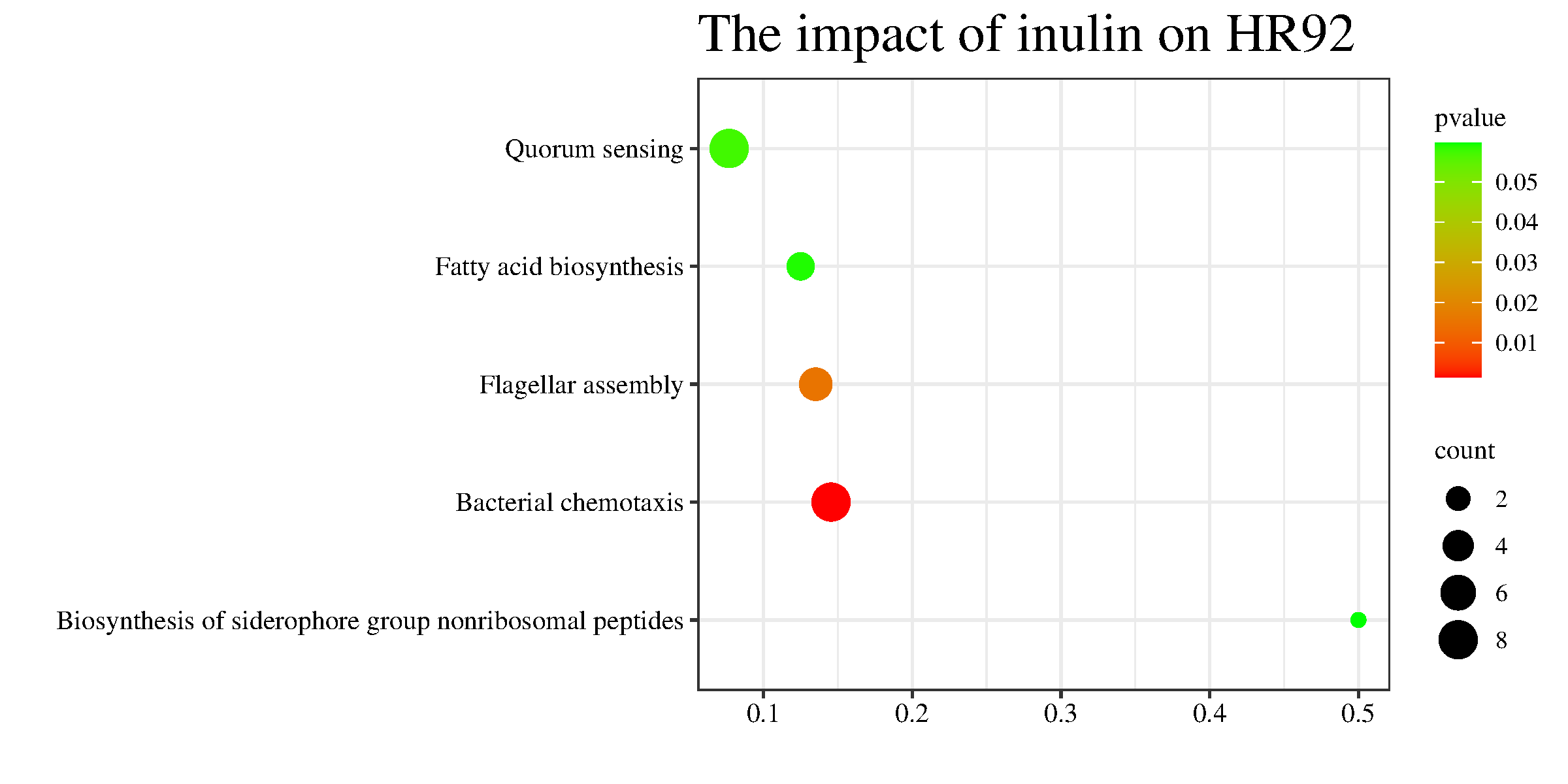

Fig. S10 Effect of inulin on HR92 gene pathways. Colors representing significance and the size of the dots indicating the number of upregulated genes

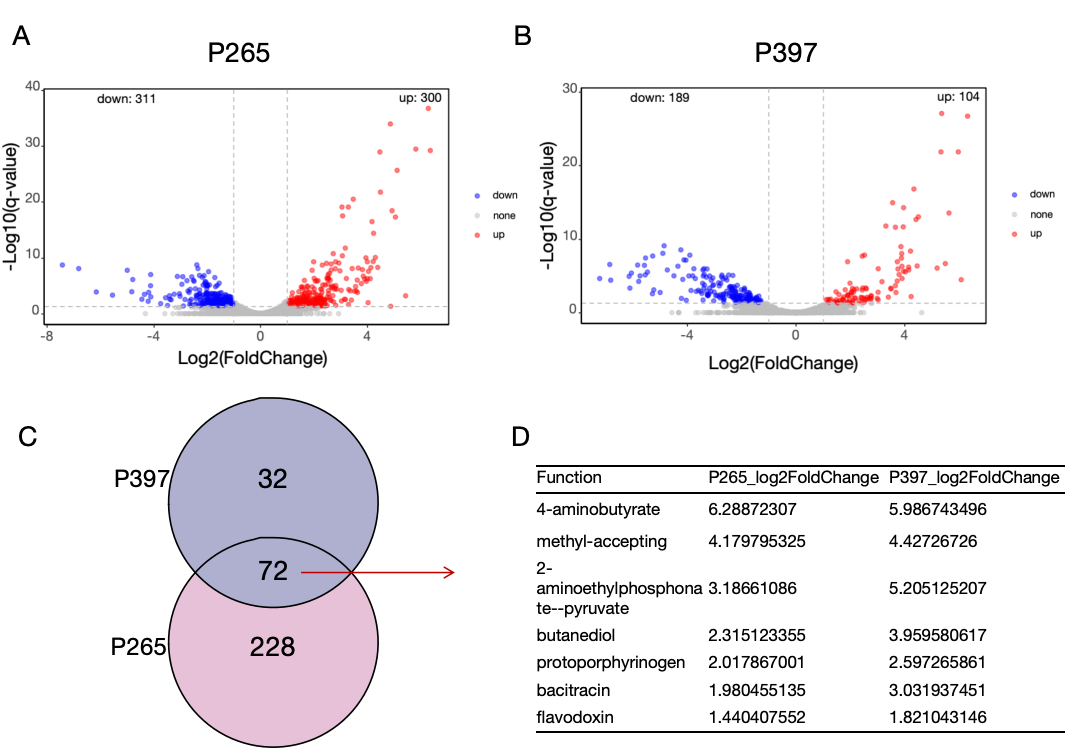

Fig. S11 Effect of *Pseudarthrobacter* spp. on HR92.

(A) Effect of P265 on HR92 gene expression, with red dots representing genes significantly upregulated and blue dots representing genes significantly downregulated (cutoff = 1). (B) Effect of P265 on HR92 gene expression, with the red dots representing genes significantly upregulated and the blue dots representing genes significantly downregulated (cutoff = 1). (C) Venn diagram showing the overlap of HR92 upregulated gene across treatments. (D) Details of genes co-upregulated by P265 and P397 in HR92.

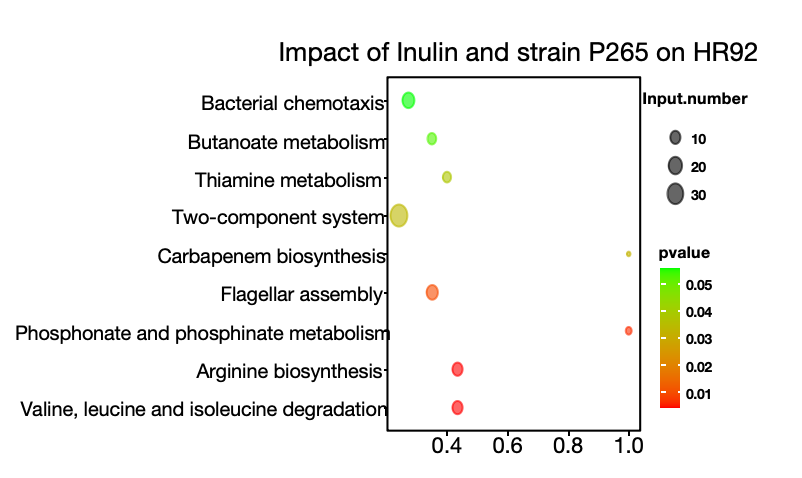

Fig. S12 Gene pathways upregulated in strain HR92 when strain P265 and inulin act together. Colors represent the level of significance and the size of the dots indicating the number of upregulated genes.

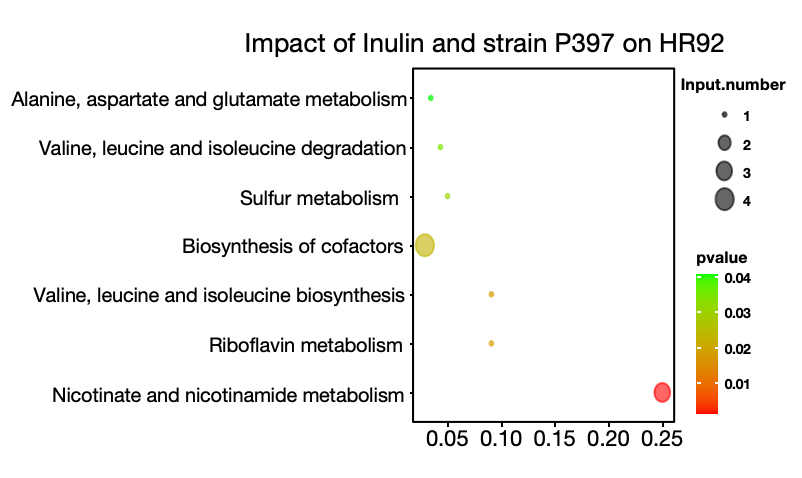

Fig. S13 Gene pathways upregulated in strain HR92 when strain P397 and inulin act together. Colors represent the level of significance and the size of the dots indicating the number of upregulated genes.

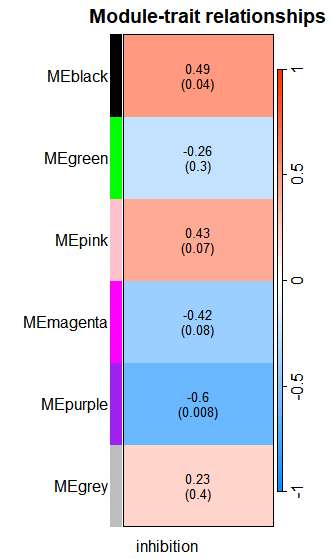

Fig. S14 WGCNA analysis of HR92 under different treatments. The heatmap colors represent the correlation between HR92 gene its ability to inhibit *R. solanacearum* , with red indicating a positive correlation and blue indicating a negative correlation. The number in the upper part of the cells indicates the specific correlation coefficient, and the number in parentheses indicates the correlation.

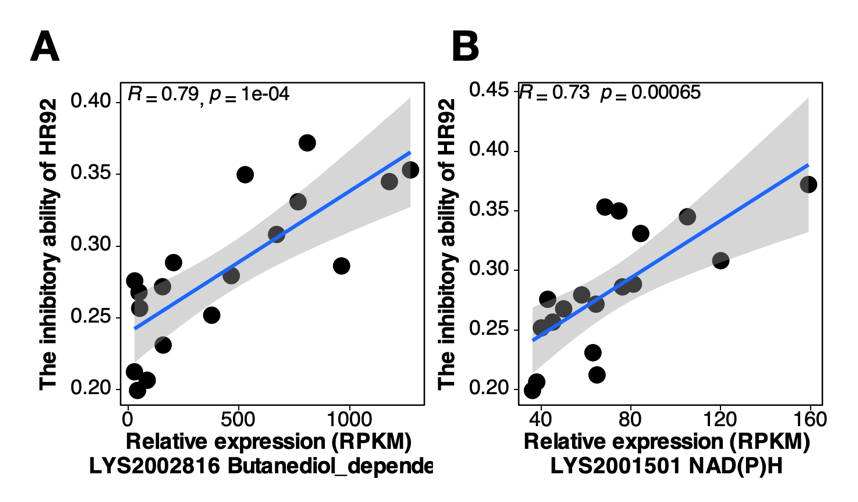

Fig. S15 Correlation between HR92 gene expression and its ability to inhibit *R. solanacearum.*

(A) Linear regression between the relative expression of butanediol dependent genes and the ability to inhibit *R. solanacearum* (*p* < 0.05). (B) Linear regression between the relative expression of NAD(P)H-related genes and the ability to inhibit *R. solanacearum* (*p* < 0.05). “R” stands for the correlation coefficient and “*p*” stands for the p-value.

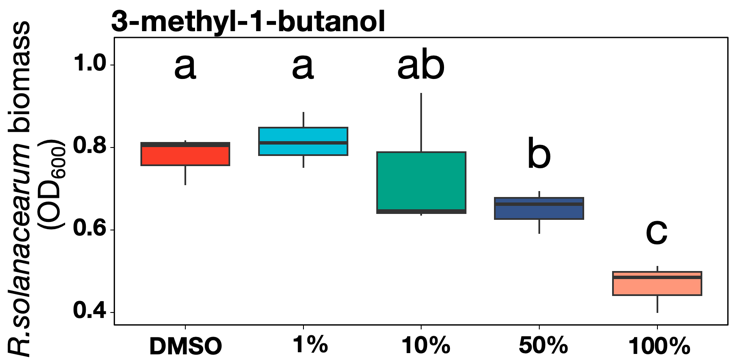

Fig. S16 Inhibition of butanedio-related volatile organic compounds against *R. solanacearum*.

Inhibition of volatile organic compound 3-methyl-1-butanol at different concentrations against *R. solanacearum*. Different letters indicate significant differences (*F_4,10_* = 6.839, *p* < 0.05, *N* = 3).

Table. S1 Potentially key bacteria genera significantly enriched in the high-concentration inulin and HR92 treatment identified using the “multipatt” package. “stat” represents the correlation coefficient.

| Genus | stat | p.value |
| --- | --- | --- |
| Pseudarthrobacter | 0.756727066 | 0.008991009 |
| Azospirillum | 0.780514898 | 0.005994006 |
| Terrabacter | 0.793884186 | 0.007992008 |
| Sandaracinus | 0.798668887 | 0.008991009 |
| Dyadobacter | 0.806206699 | 0.005994006 |
| Ellin6055 | 0.819806256 | 0.002997003 |
| Comamonas | 0.824163384 | 0.005994006 |
| Lysobacter | 0.829234262 | 0.005994006 |
| Deinococcus | 0.83978221 | 0.007992008 |
| Chondromyces | 0.841966149 | 0.002997003 |
| Variovorax | 0.854105616 | 0.002997003 |
| Cnuella | 0.868418012 | 0.002997003 |
| Pajaroellobacter | 0.871787933 | 0.005994006 |
| Agromyces | 0.88071485 | 0.002997003 |
| Qipengyuania | 0.8910551 | 0.002997003 |
| Arenimonas | 0.899258088 | 0.002997003 |
| Devosia | 0.971742719 | 0.002997003 |
| Herpetosiphon | 0.978531884 | 0.002997003 |

Table. S2 Utilization of 74 types of rhizosphere resources by P265, P397, HR92, and *R. solanacearum.* The background color represents statistical significance compared to the control. (*p* < 0.05). An asterisk (*) represents a resource utilization rate of 20% or less; Two asterisks (**) represent a resource utilization rate of 20% to 40%; Three asterisks (***) represent a resource utilization rate of 40% to 60%; Four asterisk (****) represents a resource utilization rate of 60% or more.

| resources | P265 | P397 | HR92 | RS |
| --- | --- | --- | --- | --- |
| L-Serine | ** | ** | ** | * |
| Glycine | ** | * |  | * |
| L-Homoserine |  |  |  |  |
| L-Histidine | *** | ** | ** | *** |
| D(-)-4-Hydroxyphenylglycine |  |  |  |  |
| L-Lysine | **** | *** |  |  |
| L-Alanine | *** | ** | * | *** |
| L-Leucine | ** | * | * | ** |
| L-Methionine |  |  |  |  |
| L-Phenylalanine | **** | **** | *** | **** |
| L(+)-Arginine | ** | ** | * |  |
| L-Proline | *** | ** | ** | *** |
| L-Asparagine | * | ** | * | ** |
| L-Citrulline | * | *** |  | *** |
| L-Threonine | ** | *** |  | ** |
| L-Glutamine | *** | *** | ** | **** |
| L-Tryptophan | **** | *** | *** | *** |
| L-Aspartic Acid | * | * | * | * |
| L-Ornithine | * | **** | * | ** |
| L-Valine(Val) | * |  |  | * |
| L-Pyroglutamic Acid | *** | **** | *** | **** |
| Xylitol | * | *** |  |  |
| β-Alanine | * |  |  | ** |
| D-Mannitol | **** | *** | *** | *** |
| 4-Aminobutyric Acid | *** | ** | ** | **** |
| D(+)-Trehalose Dihydrate | **** | **** | **** | **** |
| L-Isoleucine | * |  | * |  |
| Melibiose | **** | **** |  |  |
| Raffinose | **** | **** | **** |  |
| D-Mannose | **** | *** | ** |  |
| D-(+)-Cellobiose | **** | **** | ** |  |
| D(+)-Xylose | ** | **** | * |  |
| Maltose | **** | **** | **** | **** |
| D-Ribose | *** | ** |  | ** |
| β-D-Fructopyranose | **** | *** | *** | **** |
| L(+)-Rhamnose Monohydrate | * | * |  |  |
| D(+)-Glucose | **** | **** | *** | **** |
| Alpha-D-Lactose Monohydrate | **** | **** | **** | **** |
| L-Arabinose | **** | *** | ** |  |
| Calcium Pantothenate | * |  | * |  |
| D-Galactose | *** | **** | ** | *** |
| L-Ascorbic Acid |  |  |  |  |
| Nicotinic Acid |  |  |  |  |
| 3-Hydroxyflavone |  |  |  |  |
| Folic Acid |  |  |  |  |
| Quercetin |  |  |  |  |
| Riboflavin |  |  |  |  |
| Citraconic Acid |  | ** |  |  |
| Thiamine Hydrochloride | * |  |  |  |
| Shikimic Acid | *** | *** |  | *** |
| Adenine |  |  | * | * |
| Glutaric Acid | * |  |  |  |
| Nicotinamide | * | * | * |  |
| Glycolic Acid | * | * |  | * |
| Troxerutin |  |  |  |  |
| Citric Acid | ** | ** |  | *** |
| Formic Acid |  |  |  |  |
| Oxalic Acid Dihydrate |  |  |  |  |
| α-D-Galacturonic Acid |  |  |  |  |
| Fumaric Acid | * |  | * |  |
| Lactic Acid | *** | *** | ** | ** |
| Trans-Aconitic Acid |  |  |  |  |
| L-(-)-Malic Acid | ** | ** | ** | ** |
| Mucic Acid | * |  |  |  |
| 2-Ketoglutaric Acid | *** | ** | *** | *** |
| Maleic Acid |  | ** |  |  |
| Succinic Acid | ** | * | ** | *** |
| Malonic Acid | ** |  | * |  |
| D(-)-Tartaric Acid | ** | * | ** | ** |
| Pyruvicacid | * |  |  |  |
| Mucic Acid |  | * | * |  |
| Inositol |  | ** |  | **** |
| Ethanolamine |  | ** |  |  |
| Inosine | * | *** | ** |  |

Table. S3 The primers of strains

| Primers | Sequence | References |
| --- | --- | --- |
| *filc* - *R.solanacearum - Fw* | 5′-GAACGCCAACGGTGCGAACT-3′ | (46) |
| *filc* - *R.solanacearum - Rv* | 5′-GGCGGCCTTCAGGGAGGTC-3′ | (46) |
| *HR92 - Fw* | 5′-GACATCCCGTTGACCACTG-3′ | (19) |
| *HR92 - Rv* | 5′-ATTAGCTCCCTCTCGCGAG-3′ | (19) |
| 563F | 5′-GCCTCCCTCGCGCCATCAGAYTGGGYDTAAAGVG-3′ | (44) |
| 802R | 5′-GCCTTGCCAGCCCGCTCAGTACNVGGGTATCTAATCC-3′ | (44) |
| 27F | 5’-AGAGTTTGATCMTGGCTCAG-3’ | (47) |
| 1492R | 5’-TACGGYTACCTTGTTACGACTT-3’ | (47) |
